# Photoreceptor outer segment disk rim curvature relies on a tetraspanin interaction web

**DOI:** 10.64898/2026.04.29.721402

**Authors:** R. Casey Boucher, Michelle L. Milstein, Amirali Hossein, Breyanna L. Cavanaugh, Alexander J. Sodt, Andrew H. Beaven, Andrew F.X. Goldberg

## Abstract

Rod and cone photoreceptors are sensory neurons required for the first steps of image-forming vision. Mutations in *PRPH2* damage photoreceptor viability and produce a broad range of inherited retinal degenerations (IRDs) for which no treatments are available. The gene product, PRPH2 (peripherin-2), is an integral membrane tetraspanin that is required for the biogenesis and normal structure of photoreceptor outer segment (OS) disks. Peripherin-2 and its homologous partner Rom1 (rom1) self-assemble into non-covalent dimers that organize into three parallel belts that encircle each disk, forming a supramolecular scaffold that imposes extreme curvature on the disk rim. The mechanism by which curvature is generated and maintained is not yet understood. Here, we report the development of blue native (BN)-PAGE and molecular dynamics (MD) simulation approaches that suggest these proteins form an interaction web characterized by multiple interaction modes. BN-PAGE reveals that disk rim belts consist of a mixture of free peripherin-2 dimers and variously-sized chains of disulfide-linked dimers. MD simulations find that although free dimers can generate spontaneous curvature in a model membrane, dimers that are disulfide-linked into linear chains generate highly anisotropic curvature - a key hallmark of normal disk rims. The MD findings further suggest that although individual belts likely provide sufficient thermodynamic driving force to generate extreme curvature, non-covalent *inter*-belt interactions may also constrain rim diameter. The new findings advance understanding of peripherin-2 and rom1 structure and function in health and disease, and simultaneously provide promising new strategies for molecular phenotyping these clinically important proteins.

## Introduction

Peripherin-2 and rom1 are tetraspanin proteins required for the normal biogenesis and structural maintenance of vertebrate photoreceptor outer segments (OSs) (1,2). These ciliary organelles are used by rod and cone photoreceptors to transduce light into neural signals and initiate the first steps of image-forming vision (3). Tetraspanins are ubiquitous in eukaryotes, and include 33 members in humans (4). They play a broad range of biological roles but are best known as membrane domain organizers of signal transduction and trafficking processes (5).

The great majority of tetraspanins documented to date are found in numerous cell types, and participate in protein interaction webs that include both highly specific and more promiscuous associations - with themselves and with other proteins (5–7). Peripherin-2 and rom1 are unusual in this respect, given their highly photoreceptor-specific expression and nearly exclusive associations with themselves (8,9). Pathogenic mutations in the *PRPH2* gene cause a broad variety of inherited retinal degenerations (IRDs) (https://www.ncbi.nlm.nih.gov/clinvar/) which currently lack effective treatments (10,11). These diseases cause loss-of-vision and reduction in quality-of-life for patients and their families, along with economic burdens for families and societies (12). Heterozygous or homozygous loss of *PRPH2* in mice each cause profound defects for photoreceptor OS structure and retinal health (13,14). Interestingly, analogous losses of *ROM1* produce no significant, or only modest changes, to OS structure and retinal health in mouse models (15). In addition, only one definitive human case of (monogenic) *ROM1* pathogenicity has been reported to date (16). Rom1 is well-documented to interact directly with peripherin-2 (15,17,18); however, its molecular function remains to be determined.

Peripherin-2 functions at the molecular level to generate membrane curvature, an activity that is required to establish and maintain the specialized structure of both rod and cone OSs (19–21). These sensory organelles are comprised of many hundreds of membranous disks stacked one atop another, and this elaborate architecture is essential to support normal light detection by the G-protein-linked phototransduction cascade (22–24). Disks are fully (as in rods) or partially (as in cones) bounded by a high curvature (∼24nm OD) disk rim (1,25,26). Disk rim geometry represents one of the most extreme examples of membrane curvature documented in a normal biological context (27). Peripherin-2 sculpts and stabilizes these tight bends by assembling with itself and with rom1 to form a supramolecular transmembrane scaffold that generates highly negative membrane curvature (bends *away* from the cytoplasm) (21,28,29). The precise organization of this scaffold and the mechanism(s) by which it shapes OS disk rims remain to be determined.

Like other tetraspanins, peripherin-2 and rom1 assemble via non-covalent interactions between their globular extracellular 2 (EC2) domains to form stable homo- and heteromeric proteins (30,31). Originally proposed as non-covalent tetramers (32), more recent work has established non-covalent dimers as the minimal structural unit (29). Importantly, and unlike other tetraspanins, peripherin-2 and rom1 EC2 domains possess a free cysteine (C150 and C153 respectively) that mediate the assembly of disulfide-linked “higher-order complexes” (33,34). These complexes have been resolved as non-covalent dimers linked into chains of varied lengths by disulfide bonds (21,29), and cryo-electron tomography data suggests that three parallel rows (referred to here as “belts”) of peripherin-2/rom1 polymers encircle disks to form a transmembrane scaffold (28). Figure 1A provides a schematic overview of protein organization at the disk rim.

**Figure 1.**
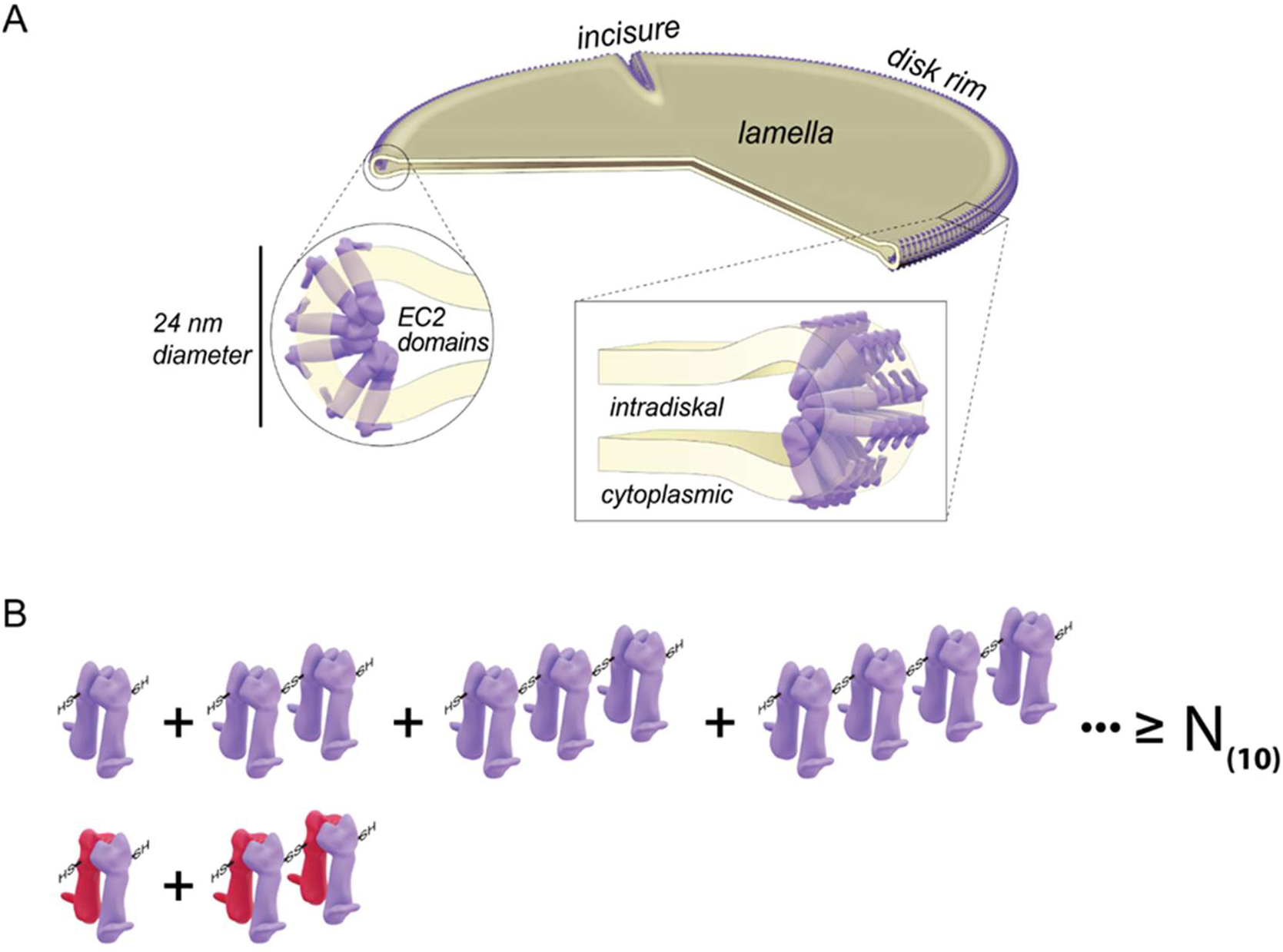
OS disk rim scaffolding by peripherin-2 and rom1 polymeric forms. (A) OS disk (*yellow*) showing the localization and organization of peripherin-2 (purple) at the rim. Three parallel polymer belts of peripherin-2 encircle the disk perimeter. Detailed views of the extreme rim curvature illustrate V-shapes within individual non-covalent dimers and between the central and peripheral dimer belts that form the supramolecular scaffold (21,28,29). Although rom1 (not illustrated) is normally present within this structure, its precise organization is not known. Rom1 is not required to generate normal OS disk rims (42). (B) Current model for polymeric forms of peripherin-2 and rom1 (*red*) solubilized from OS membranes and transfected HEK293 cells using gentle detergents. Minimal structures include non-covalent homo- and hetero-typic dimers. Covalent assembly of non-covalent dimers into variously sized disulfide-linked chains is mediated by C150 (peripherin-2) and C153 (rom1) residues within the EC2 domains (21,29,33,34). Rom1 is confined to dimeric and tetrameric forms.

To achieve the goal of understanding *PRPH2* genotype-phenotype relationships, a deeper knowledge of peripherin-2/rom1 organization within the disk rim scaffold, combined with a mechanistic understanding of the means by which it generates membrane curvature is needed. The studies described here address these questions by, 1) providing new methods for assessing mutational impacts on peripherin-2 self-assembly, 2) documenting a web of protein-protein interactions that support the disk rim scaffold, and 3) identifying disulfide-linked chains of non-covalent peripherin-2 dimers as key structures for generating the anisotropic spontaneous membrane curvature that constrains disk rim shape and diameter.

## Results

### A reoptimized velocity sedimentation method improves assay reproducibility and dynamic range and emphasizes polymer heterogeneity

Peripherin-2 and rom1 are detergent-extracted from rod outer segment (ROS) disk membranes (and transfected HEK293 cells) as linear polymers of varied lengths (21,29). Because the precise organization of these two proteins within the disk rim scaffold remains undefined (28), we reoptimized a commonly used velocity sedimentation method (2,18,34) to gain additional insight into peripherin-2 and rom1 structure. Reproducibility was improved by developing a semi-automated method to fractionate small-volume sucrose density gradients (Figs. S1, S2), and dynamic range was expanded by reducing sedimentation g-force. We applied the reoptimized method to compare protein behavior in the two detergents commonly used to study these proteins.

Representative velocity sedimentation results are presented in Figure 2 (A, B). Bovine ROS membrane extracts, solubilized with either Triton X-100 (TX) or dodecylmaltoside (DDM), were sedimented and fractionated, and the fractions were analyzed by non-reducing SDS-PAGE followed by western blotting. Importantly, both peripherin-2 and rom1 were retained within the sucrose gradients, regardless of solubilizing detergent, and no pelleting of either protein was observed. This contrasts with the original sedimentation method, in which a substantial portion of the peripherin-2 is found in the pellet fraction (21). Consistent with previous studies, SDS-denatured peripherin-2 and rom1 are each observed as monomeric and disulfide-linked dimeric (SS-D) forms by non-reducing SDS-PAGE (35). SS-D forms are not native disk rim structural elements, but are only released upon SDS denaturation of disulfide-linked chains of non-covalent dimers during analysis (21,34). Under the reoptimized conditions, Triton X-100-solubilized peripherin-2 was distributed in a single broad peak throughout all gradient fractions (Fig. 2A, *top panel*), consistent with the original demonstration that it is solubilized as heterogeneously-sized complexes (34). DDM-solubilized peripherin-2 was similarly distributed (Fig. 2B, *top panel*), though it sedimented slightly faster (Fig. 2C, *top panel*). In contrast, rom1 was restricted to smaller complexes (lighter gradient fractions), regardless of which detergent was used for solubilization (Fig. 2A, B, *middle panels*). Like peripherin-2, rom1 mobility increased slightly when solubilized with DDM vs. Triton X-100 (Fig. 2C). Overall, data from the reoptimized sedimentation approach is consistent with previous findings, but the modified approach offers improved dynamic range, reproducibility, and ease-of-use. The results emphasize that detergent-solubilized peripherin-2 complexes from ROS membranes sediment in a very broad manner, suggesting a continuum of numerous polymeric forms.

**Figure 2.**
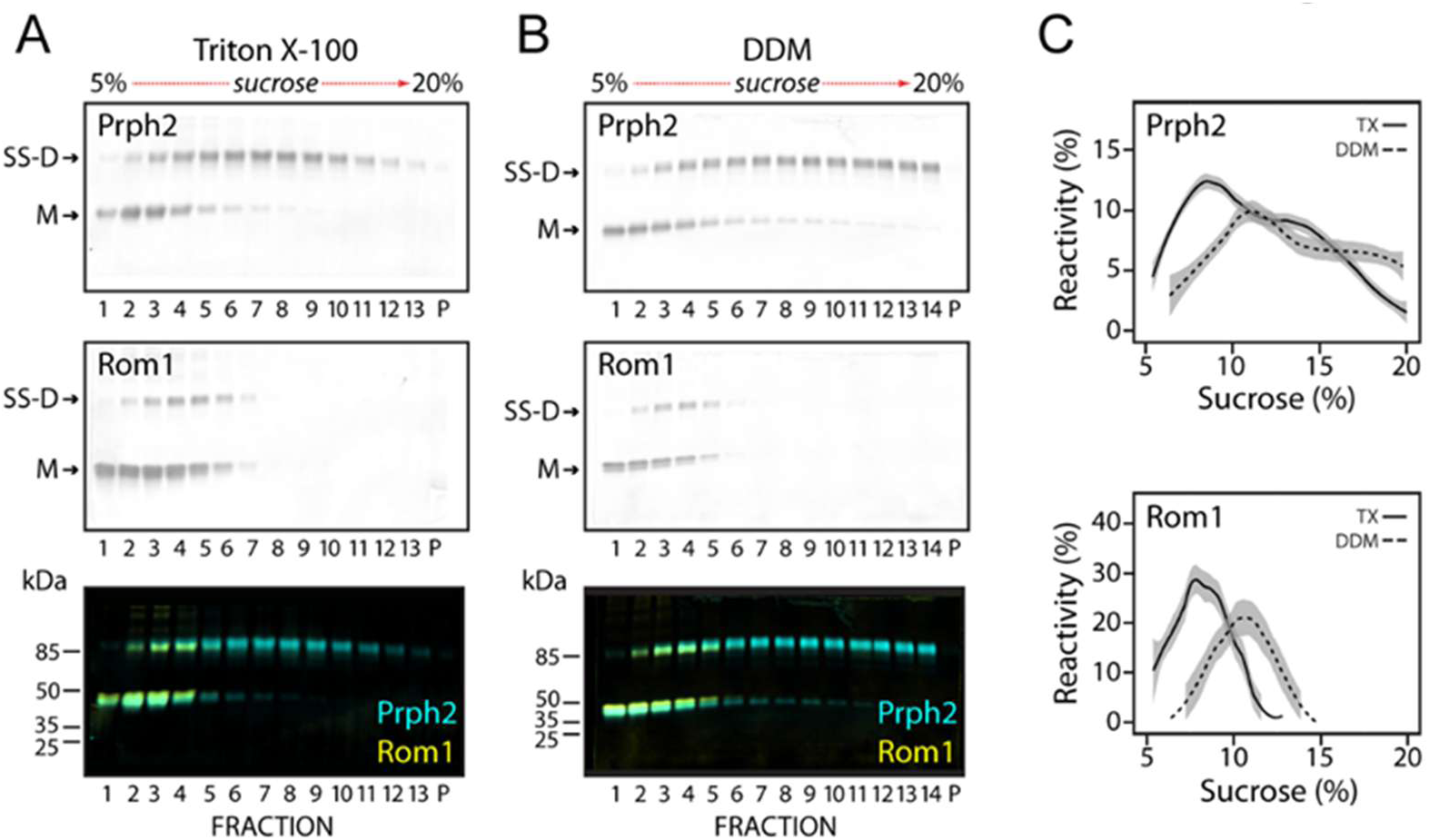
Reoptimized velocity sedimentation analysis demonstrates a broad heterogeneity of peripherin-2 and rom1 polymeric forms. Detergent extracts of bovine ROS membranes (solubilized with either 1% Triton X-100 or 1% DDM) were subjected to velocity sedimentation under optimized non-reducing (and non-denaturing) conditions on 5 to 20% sucrose gradients. Gradient fractions were analyzed by non-reducing SDS-PAGE western blots uing anti-peripherin-2 (Prph2) and anti-rom1 (Rom1) antibodies. Prph2 and Rom1 each migrate as monomeric (M) and disulfide-linked dimeric (SS-D) species. SS-Ds are not autonomous disk rim structural elements, but are released by SDS denaturation of disulfide-linked chains of non-covalent dimers during analysis (21,34). (A) Western blot showing sedimentation profiles for Triton X-100 solubilized Prph2 (*top panel*), Rom1 (*middle panel*), and merged data (*bottom panel*). (B) Western blot showing sedimentation profiles for DDM solubilized Prph2 (*top panel*), Rom1 (*middle panel*), and merged data (*bottom panel*). C) LOESS curves comparing sedimentation profiles in Triton X-100 (n=11) and in DDM (n=3) for Prph2 (*top panel*) and Rom 1 (*bottom panel*). Shading indicates 95% confidence levels.

### Blue Native-Polyacrylamide Gel Electrophoresis (BN-PAGE) resolves peripherin-2 and rom1 polymer chain lengths and relative abundance

Early studies of bovine peripherin-2 and rom1 observed that they assemble into tightly associated non-covalent homomeric and heteromeric complexes (18), and that disulfide-mediated oligomerization of “non-covalent core complexes” (via C150 in peripherin-2 and C153 in rom1) creates larger species (34,36). Recent investigations conclude that the larger species described by those early studies are variously-sized polymers of noncovalent dimers linked into linear chains by disulfide bonds (21,29). To date however, an ensemble method to determine chain lengths and relative abundance has not been available.

To address this knowledge gap, we developed a BN-PAGE approach for resolving detergent-solubilized peripherin-2 and rom1 polymer chains by length. Bovine ROS membranes were solubilized with either Triton X-100 or DDM, and proteins were analyzed by BN-PAGE and western blotting. In each detergent, peripherin-2 generated laddered banding patterns spanning a broad range of polymer sizes (1 to >10 dimers), while rom1 was only present in smaller polymers, primarily dimers and tetramers (Fig. 3, A-C). For each protein, the smallest polymers were the most abundant (Fig. S3). These bands were identified as non-covalent dimers by reference to the minimal structural unit resolved by single particle cryo-electron microscopy (29). The non-covalent dimer migrated at an apparent molecular weight (MW) of ∼100 kDa by BN-PAGE relative to NativeMark standards. This is ∼25 kDa larger than that predicted for a peripherin-2/rom1 heterodimer, likely reflecting the detergent micelle contribution (37). Triton X-100 solubilization produced the most highly resolved ladders, and the reduction of Trion X-100 extracts with dithiothreitol (DTT) prior to BN-PAGE reproducibly collapsed larger polymers into smaller (1-3 dimer) species (Fig. 3A). BN-PAGE analyses of DDM solubilized ROS produced ladders similar to those seen with Triton X-100, but they included additional minor intermediate bands. A plot of peripherin-2 apparent MW as a function of predicted N-mer shows a linear relationship, regardless of solubilizing detergent (Fig. 3D). The results suggest that polymer lengths increase by a constant increment corresponding to the non-covalent dimer. In sum, the BN-PAGE data indicate that an ensemble of stable species, ranging from single non-covalent dimers to polymer chains of >10 disulfide-linked dimers, were solubilized from native ROS membranes, with rom1 being restricted to dimeric and tetrameric forms (illustrated in Fig. 1B).

**Figure 3:**
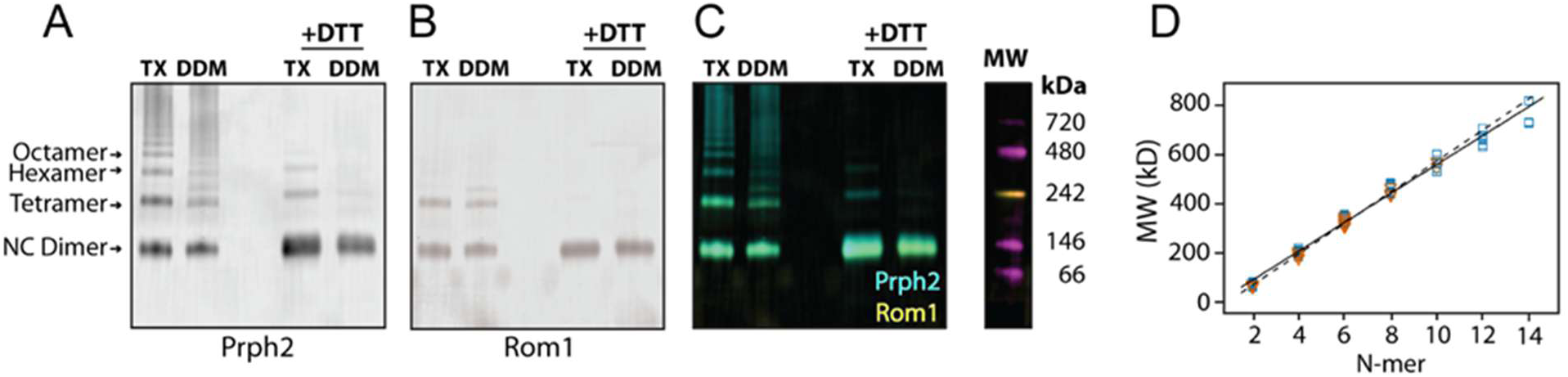
A BN-PAGE approach resolves detergent-solubilized Prph2 and Rom1 polymers by size. (A-C) Proteins were extracted from native ROS membranes with either 1% Triton X-100 (TX) or with 1% DDM, then were analyzed by non-reducing BN-PAGE western blotting for Prph2 and Rom1. (A) Prph2 shows laddered banding patterns spanning a broad range of polymer sizes in each detergent, from 1 to ≥ 10 noncovalent dimers. DDM solubilization often generated additional intermediate bands superimposed on the ladders seen in Triton X-100. In each detergent, reduction of samples prior to BN-PAGE (+DTT) collapsed ladders to mainly dimers. (B) Rom1 shows a relative absence of laddering compared to that seen for peripherin-2, with major bands confined to dimer and tetramer species. In each detergent, reduction of samples prior to BN-PAGE (+DTT) converted tetramers to dimers. (C) Merged Prph2 and Rom1 data. (D) A plot of apparent MW as a function of N-mer size for Prph2 in Triton X-100 (*blue squares*) and DDM (*orange triangles*) shows a direct relationship. Data includes replicate lanes from 3 independent blots (total n = 11 lanes for Triton X-100 and n = 12 lanes for DDM). Statistical regression shows a strong linear correlation in both Triton X-100 (solid line, R^2^ = 0.985, P < 0.001) and DDM (dashed line, R^2^ = 0.997, P < 0.001).

### BN-PAGE analysis of polymer lengths in ROS sedimentation fractions

Since its first implementation (18), velocity sedimentation has been used in numerous studies to assess peripherin-2/rom1 complex formation (2,12). Although this technique continues to be used to probe the impact of *PRPH2* mutations - its utility is limited by low throughput and the inability to report on species stoichiometry and abundance. We therefore combined our reoptimized sedimentation procedure with the new BN-PAGE method to evaluate the resolving power of the sedimentation approach.

Figure 4 (A, B) presents BN-PAGE western blot data for sedimented and fractionated peripherin-2 and rom1 solubilized from bovine ROS membranes. The blots show stair-like banding patterns with increasing sucrose gradient density, regardless of whether membranes were solubilized with Triton X-100 or DDM. Peripherin-2 was observed in a broad range of polymer chain sizes (1 to >10 dimers), while rom1 was only present in smaller polymers (1 or 2 dimers). Plots of peripherin-2 and rom1 polymer chain abundance in the fractionated gradients illustrate that peaks for individual species show considerable overlap (Fig. 4C, D). These data demonstrate that although velocity sedimentation fractionates polymers by size, it does not resolve or identify individual species. In contrast, BN-PAGE provides a more informative approach, enabling resolution and quantification of the length and relative abundance of numerous peripherin-2 and rom1 polymer species.

**Figure 4:**
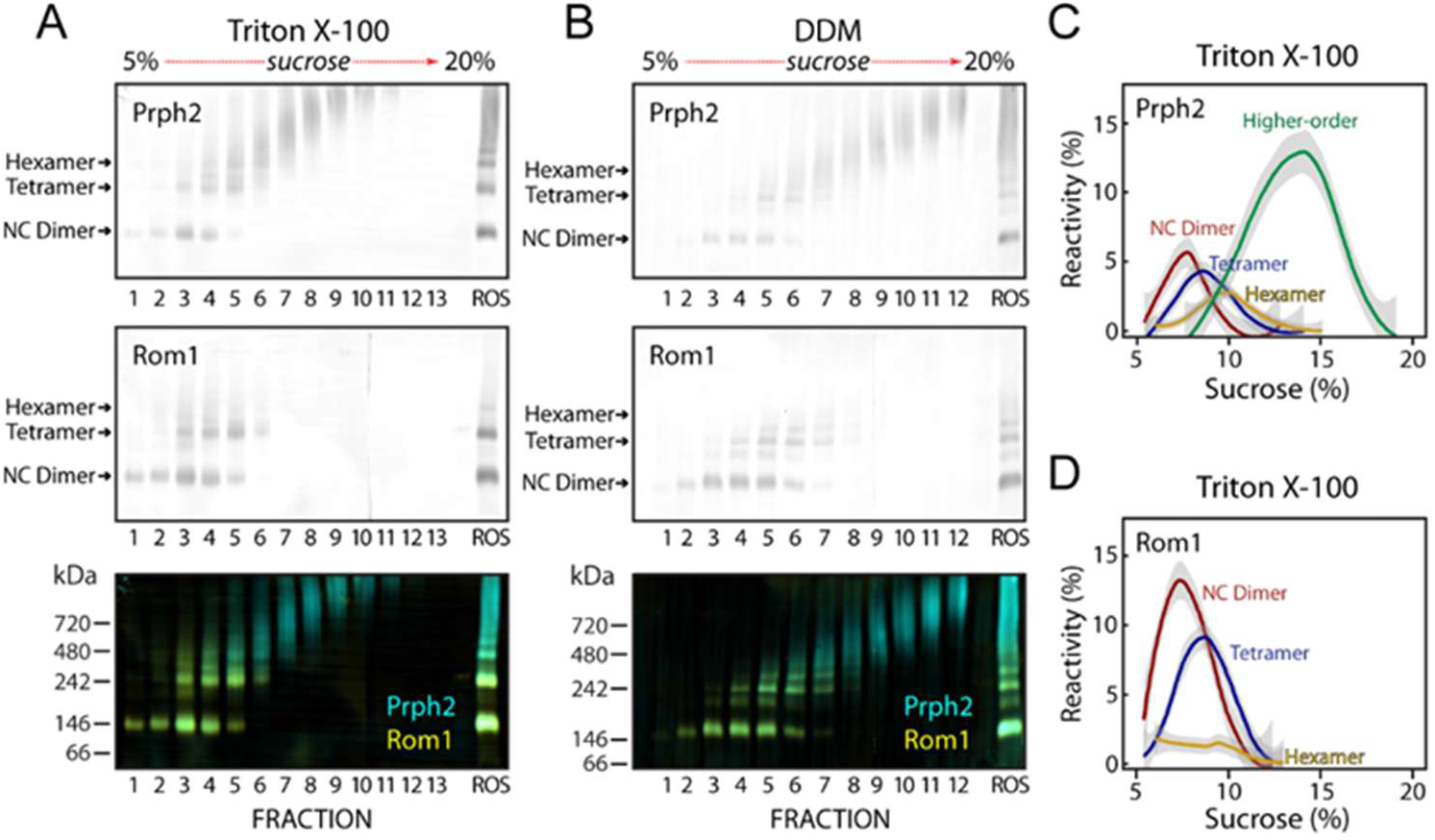
BN-PAGE of sedimented and fractionated ROS lysates reveals limitations of velocity sedimentation. Purified bovine ROS membranes were solubilized with (A) 1% Triton X-100 or with (B) 1% DDM, and then extracts were sedimented and fractionated as in Fig. 2. Gradient fractions were analyzed by non-reducing BN-PAGE western blotting; Prph2 (*top*), Rom1 (*center*), and merged (*bottom*) data channels are shown for each detergent. Prph2 polymers range from 1 to ≥10 non-covalent dimers, while Rom1 lacks higher-order species and is largely confined to dimers and tetramers. Plots comparing sedimentation profiles in Triton X-100 show relative abundance of (C) Prph2-containing and (D) and Rom1-containing species. Lines represent LOESS regressions of relative abundances for each size species indicated (*Red: NC Dimer; blue: tetramer; yellow: hexamer; green: higher-order*). Data represent 8 independent sedimentation trials using Triton X-100 solubilized material. Grey shading illustrates 95% confidence intervals.

### BN-PAGE molecular phenotyping reveals mutation-induced *PRPH2* self-assembly defects

*PRPH2* mutations cause a broad variety of IRDs with high variability in expressivity and penetrance (10). Because the new BN-PAGE method proved useful for analyzing peripherin-2 and rom1 from ROS membranes, we next assessed its efficacy for screening mutant protein variants expressed in HEK293 cells. This platform has been a reliable tool for understanding disease-associated *PRPH2* mutations (38), and we sought to expand its utility by combining it with BN-PAGE analysis of *PRPH2* variant self-assembly.

Figure 5A shows a BN-PAGE analysis comparing HEK293-expressed mutant and wild-type (WT) peripherin-2 variants solubilized in Triton X-100 or DDM. In each detergent, WT peripherin-2 banding patterns recapitulated those seen for ROS membranes (Figs. 3, 4). Recombinant WT peripherin-2 produced laddered banding patterns spanning a variety of polymer sizes, ranging from one to >10 dimers. This was anticipated, because prior work has shown that rom1 is not required for recombinant peripherin-2 self-assembly in HEK293 or COS cells (18,34,38). In contrast, a C150S mutant produced no laddering, but instead just one major band that migrated as a dimer, and one minor band that migrated as a tetramer. This result reflects the known dependence of disulfide-linked peripherin-2 dimer chains on C150, and suggests that non-covalent dimers may interact with themselves in a non-covalent manner to form higher-order forms. We next examined a pathogenic L185P mutation associated with retinitis pigmentosa (39,40). Like C150S, the L185P variant also produced a single major band, which migrated as a dimer of slightly reduced mobility (Fig. 5A). This finding is consistent with evidence that L185P prevents normal subunit assembly (41), and traps the protein in an abnormally oxidized form (21). We hypothesized that L185P exerts this effect by disrupting the EC2 domain dimerization interface. We tested this idea using a C150S;L185P double mutant and predicted that this combination would generate a monomeric protein. BN-PAGE analysis of the C150S;L185P mutant shows a predominant band that migrated as a (∼50 kDa) monomer. To independently verify the BN-PAGE results, the reoptimized sedimentation method was applied to the WT, C150S, and C150S;L185P variants (Fig. 5B-G). L185P sedimentation has been published previously (21). In each case, the sedimentation velocity findings are in agreement with the BN-PAGE analyses. Together, the results demonstrate that BN-PAGE offers a robust approach for efficiently screening *PRPH2* mutations for defects in non-covalent dimerization and disulfide-mediated dimer.

**Figure 5:**
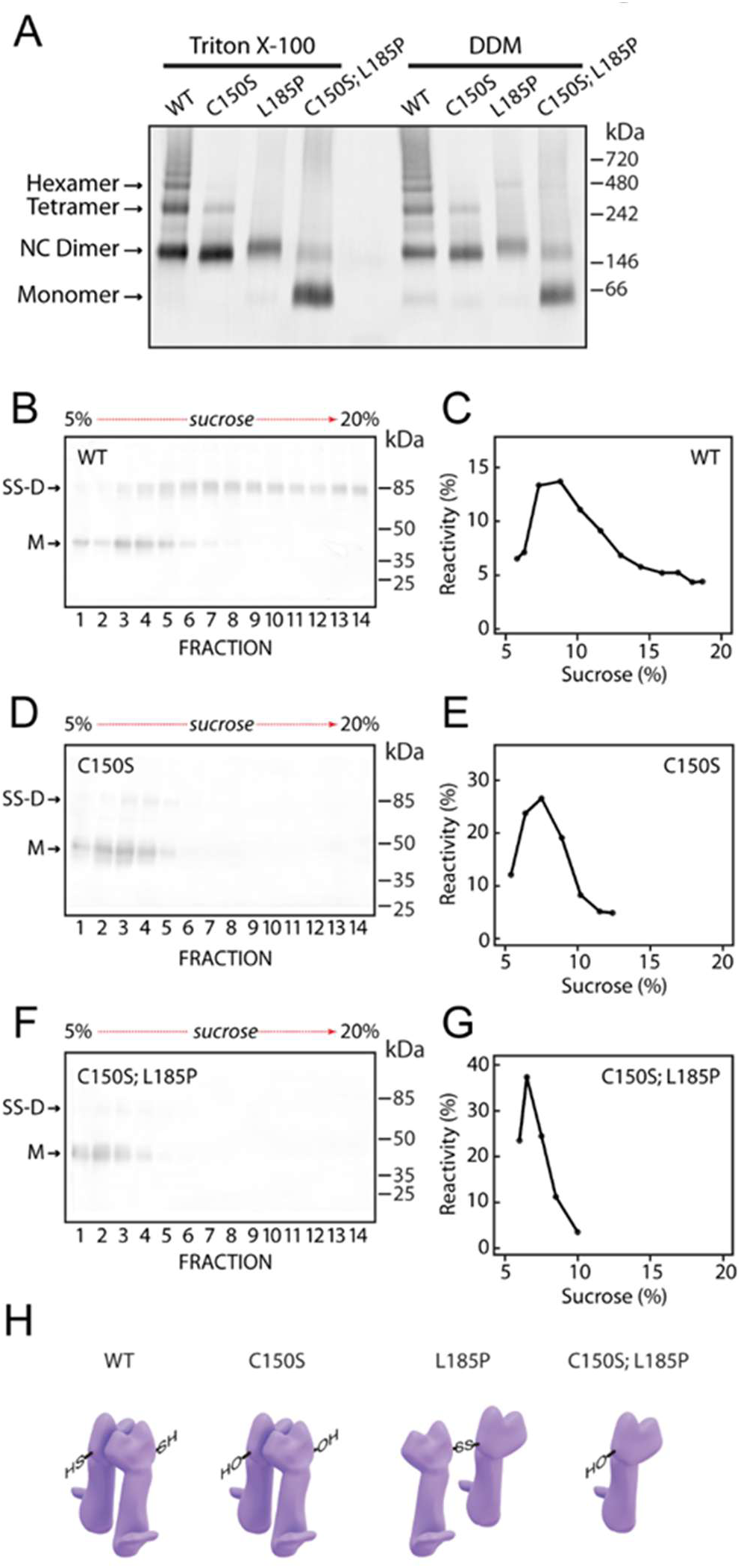
BN-PAGE offers an effective method for screening recombinant peripherin-2 variants. (A) Prph2 variants were expressed in HEK293 cells, extracted into either 1% Triton X-100 (left lanes) or 1% DDM (right lanes), and then were analyzed by BN-PAGE followed by anti-Prph2 western blotting. Similar results were observed with each detergent. The previously studied C150S mutant, which lacks the key cysteine required for *inter*molecular disulfide formation (33), formed non-covalent (NC) dimers, and a minor tetramer band, but no higher-order polymers. The well-documented L185P mutant disrupted non-covalent Prph2 dimerization and instead formed improperly oxidized dimers (21), which blocked higher-order polymer formation. A novel C150S;L185P double mutant, in which non-covalent dimerization and intermolecular disulfide formation were both simultaneously blocked, produced a predominantly monomeric form of the protein. BN-PAGE results were confirmed using non-reducing sedimentation analyses and profiles for: WT (B, C), C150S (D, E), and C150S;L185P (F, G) mutants expressed in HEK293 and solubilized with Triton X-100. *M, monomeric*; *SS-D, disulfide-linked dimeric*. (H) Physical models for Prph2 mutants: WT, showing the NC dimer form (larger polymers are omitted for clarity); C150S, a NC dimer incapable of disulfide-chain formation, L185P, a pathogenic mutation that inhibits normal NC dimerization and forms improperly oxidized dimers; C150S;L185P, a double mutant that traps the protein as a monomer.

### BN-PAGE identifies defective peripherin-2 polymerization in a mouse model of *Rom1*-associated IRD

Peripherin-2 and rom1 normally interact and work together to generate and maintain OS structure, but they are not of equal importance. Peripherin-2 is essential for scaffolding disk rim structure and OS organelle biogenesis - photoreceptors in *Prph2* null (*rds* homozygous) mice fail to produce OS organelles. In contrast, rom1 molecular function remains to be defined and is not essential for OS formation. Photoreceptors in *Rom1* knockout (KO) mice develop OSs with only mild structural anomalies (13–15,42). We used *Rom1* KO mice to investigate rom1 contribution to peripherin-2 polymerization, and simultaneously evaluate BN-PAGE as a general tool for molecular phenotyping of *PRPH2*-associated mouse IRD models.

ROS membranes were purified from *Rom1* KO and from WT mice, and non-reducing SDS-PAGE was used to compare the peripherin-2 content from each under denaturing (but non-reducing) conditions (Fig. 6A). In each case, two major peripherin-2 bands were observed on western blots, corresponding to monomeric and disulfide-linked dimer (SS-D) species. As previously mentioned, SS-D forms are not native disk rim structural elements, but are released by SDS denaturation of disulfide-linked chains of non-covalent dimers during analysis. Comparison of ROS purified from multiple animals showed that monomer to SS-D ratios did not vary between individuals, nor were they altered by the absence of rom1. BN-PAGE analysis of these same samples showed that rom1 absence created a substantial loss of the highest-order peripherin-2 polymer species (Fig 6B). To test this result using an independent method, the samples were also analyzed by the reoptimized velocity sedimentation method (Fig. 6C). Here again, the absence of rom1 (*top* vs *bottom panel*) substantially reduced the abundance of higher-order peripherin-2 polymers. Finally, fractionated gradients from the sedimented *Rom1* KO and WT ROS samples were analyzed by BN-PAGE (Fig. 6D). Consistent with the sedimentation results, BN-PAGE showed a redistribution of peripherin-2 from larger-to-smaller polymeric forms when rom1 was not present (Fig. 6E). Overall, the findings demonstrate that rom1 supports the assembly of the largest peripherin-2 polymers, and that BN-PAGE offers an effective method for molecular phenotyping of peripherin-2 in IRD model mice.

**Figure 6:**
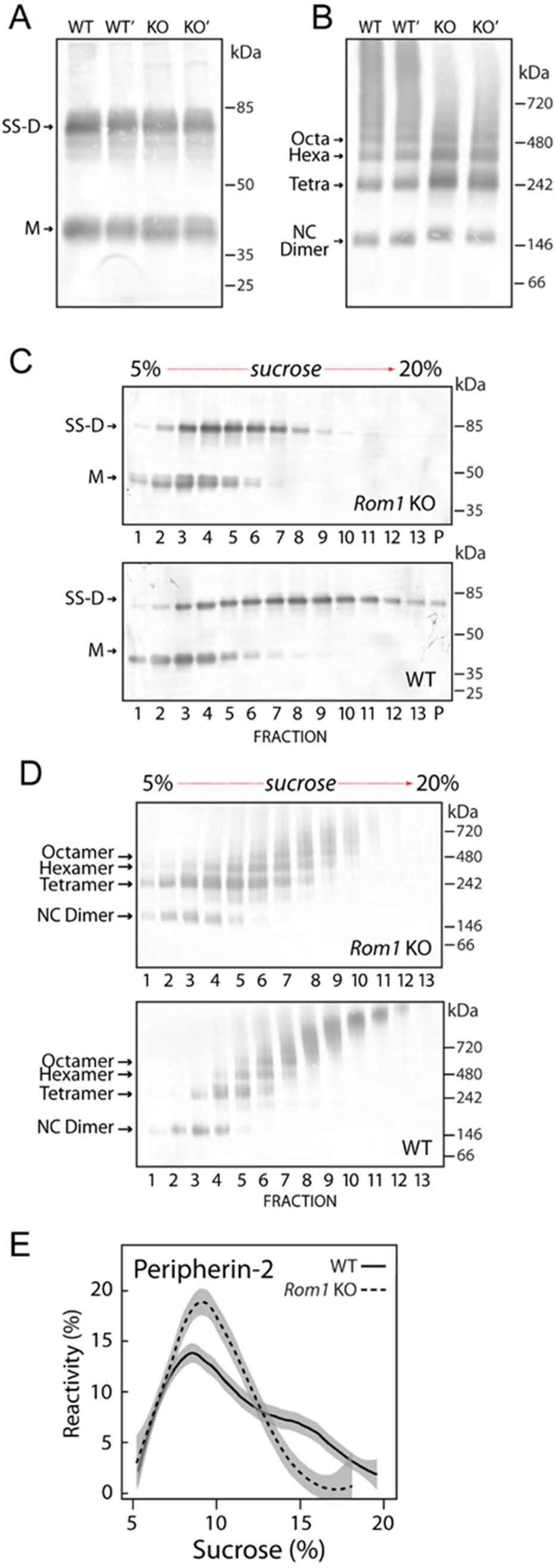
BN-PAGE vs. velocity sedimentation as phenotyping tools for characterizing peripherin-2 polymerization in a murine IRD model. (A) Non-reducing SDS-PAGE western blotting, using independent purifications of ROS membranes from two wildtype (WT, WT’) and two *Rom1* knockout (KO, KO’) mice. (B) BN-PAGE western blotting of Triton X-100 ROS extracts from two WT and two *Rom1* KO mice. A loss of the highest-order peripherin-2 species is seen in the *Rom1* KOs. (C) Sedimentation analysis of Prph2 in Triton X-100 solubilized ROS membranes purified from *Rom*1 KO (*upper panel*) and WT (*lower panel*) mice (M, monomeric; SS-D, disulfide-linked dimeric) by non-reducing SDS-PAGE western blotting. A loss of the highest-order Prph2 species is seen in the *Rom1* KO. (D) BN-PAGE western blotting of the sedimentation fractions from *Rom1* KO (*upper panel*) and WT (*lower panel*) mice shown in *panel C*. Gradients from *Rom1* KO mice show a loss of highest-order and increase in mid-sized polymers relative to WT. (E) Plots show sedimentation profiles for Prph2 present in ROS extracts from WT (*solid line*) and *Rom1* KO (*dashed line*) mice. Lines represent LOESS regressions from collations from ten WT and eight *Rom1* KO independent sedimentation runs; grey shading illustrates 95% confidence intervals.

### Molecular modeling predicts that non-covalent interactions can organize peripherin-2 homo and heteromeric dimers into linear polymers

Evidence from multiple laboratories indicates that non-covalent peripherin-2 dimers are extracted from ROS membranes (and transfected HEK293 cells) as disulfide-bonded chains of varied lengths (21,29). Given the observation that non-chain forming C150S mutant dimers can weakly self-associate (Fig. 5A), we hypothesized that WT dimers may also interact non-covalently and used molecular modeling to evaluate this idea.

AlphaFold2-Multimer (AF-M) was utilized to identify potential interactions between peripherin-2 homodimers. The top-ranked model predicts a tetrameric complex consisting of two side-by-side homodimers, which form a linear polymer along the plane formed by the non-covalent dimerization interface (Fig.7B). This organization mirrors the linear structure of peripherin-2-containing polymers revealed by transmission electron microscopy (TEM) (21), single-particle cryo-EM analysis (29), and cryo-ET (28). The dimer-dimer interaction interface is mediated entirely by the EC2 domains, and appears to include close contacts between intramolecular salt bridges (D145/R183) located on opposing (and diagonally situated) monomers of the neighboring dimers (Fig. 7C). Significantly, this interaction also brings the C150 residues from these same two monomers into close proximity.

**Figure 7:**
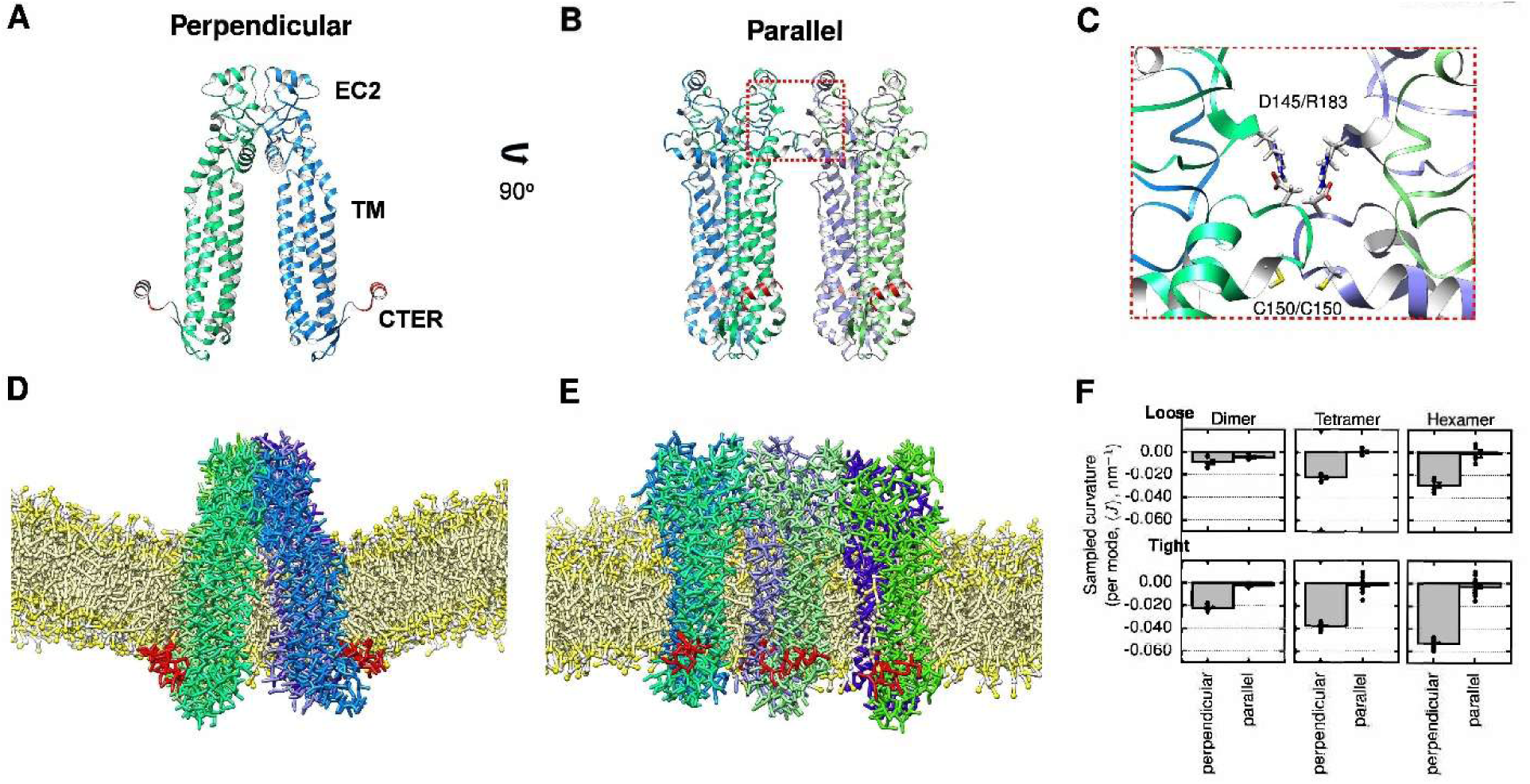
Molecular dynamics simulations predict the generation of highly anisotropic membrane curvature by peripherin-2 chains. AlphaFold Multimer predicted (A) non-covalent dimer and (B) non-covalent tetramer structures. The dimer prediction bears close similarity to the experimentally determined structure (29). (C) The tetramer prediction suggests that close contacts between intramolecular salt bridges (R183/D145) located on opposing (and diagonally situated) monomers of the neighboring dimers can help stabilize non-covalent dimer-dimer interactions that bring free C150 residues into close proximity. (D, E) Representative images from MD simulations are shown for a hexamer chain model. Individual chains within non-covalent dimers are represented colored green and blue, C-terminal amphipathic helixes are illustrated in red, and lipids are given in yellow. Water and ions are not shown for clarity. (F) Curvature sampled per mode by the dimeric, tetrameric, and hexameric peripherin-2 complexes (0.0025, 0.0050, and 0.0075 dimers/nm^2^ surface coverage respectively) is plotted for loosely (*top*) and tightly (*bottom*) restrained protein models.

### Molecular dynamics simulations suggest that an interaction web underlies peripherin-2 generation of extreme curvature

Despite significant recent advances for understanding peripherin-2 structure and function, it is not yet clear how the energetically unfavorable curvature of the disk rim is stabilized. The intrinsic elastic energy of biological lipid bilayers requires the input of substantial energy to achieve curvatures with outer diameters of 24 nm (27,43). Evidence to date suggests that the vast majority, or perhaps all, of that energy can be supplied by peripherin-2 homotypic interactions (15). According to the Helfrich/Canham elasticity model of bilayer curvature (43,44),

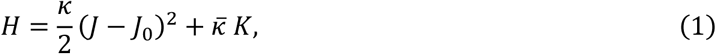

∼0.15 kT of energy is required to bend a 1 nm^2^ membrane patch into the approximate radius *R*=10 nm of the disk rim (20 nm diameter). For modeling purposes, curvature is measured at the bilayer *midplane*, as opposed to the outer edge of the bilayer that is resolved by TEM. At the midplane area (ca. 65 nm^2^ per dimer) ∼10 kT is required. For this estimate, 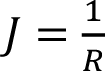 is the curvature of the disk rim, 𝐽_0_ is the spontaneous curvature (here assumed to be zero for a symmetric bilayer initially free of protein), 𝜅 ≈ 30 kT is the bending modulus (the force constant for variations in curvature, which can vary substantially with conditions (45), and 𝜅̅ is the modulus of Gaussian curvature (for the rim this term is neglected; see below). Molecular modeling translates the interactions between protein shape and lipids into a curved membrane deformation, implying a change in 𝐽_0_ - potentially making disk rim curvature spontaneous. To understand how the extreme curvature of a disk rim is stabilized and to what extent pathogenic *PRPH2* mutations may impair protein activity for curvature generation, it is necessary to elucidate these interactions in detail, and determine how they couple, to yield 𝐽_0_ values sufficient to spontaneously create disk rim curvature.

Here, we used a Martini coarse-grained model (46) to investigate the extent to which peripherin-2 homodimers, both singly and in individual disulfide-linked chains, could contribute to the extreme curvature of OS disk rims by altering the spontaneous curvature of a bilayer patch. The Martini model was selected because, 1) its lipid bilayer model has curvature energetics closely matching experiment and detailed “all-atom” models (47), and 2) protein shape, including that of transmembrane and amphipathic alpha helices, is well above this model’s level of coarseness (48). Because Martini protein models often exhibit tertiary structure instability and a tendency to unfold, they are typically maintained near a target conformation using restraints (49). We therefore compare two common approaches here. A *loosely* restrained model that defines secondary structure elements without external control of tertiary structure, and a *tightly* restrained model with fixed tertiary structure that maintains the AlphaFold-derived initial condition.

Simulated patch spontaneous curvature 𝐽_0,patch_ is computed by the area-weighted fraction of protein spontaneous curvature and that of the bilayer (here assumed to be of symmetric composition, and thus of spontaneous curvature zero). The key model parameter is thus the *protein* spontaneous curvature 𝐽_0,protein_, which is estimated from the curvature sampled by a protein per independently-fluctuating normal mode, applying theory developed by Sapp, et al (50):

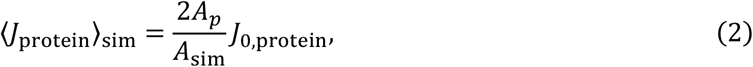

where here 𝐴_p_ is the projected area of the protein complex (here doubled because the protein affects both the upper and lower leaflets), 𝐴_sim_ is the area of the simulation, and 〈𝐽_protein_〉 is the curvature sampled by the protein averaged over the trajectories and replicates, measured at the geometric center of the complex. Assuming the surrounding bilayer has zero spontaneous curvature, the bilayer spontaneous curvature resulting from the protein contribution at target (disk rim) coverage is modeled according to:

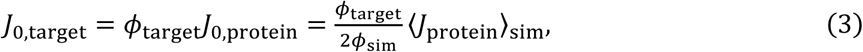

where 𝜙_target_ is the target fractional coverage of the bilayer, with 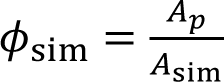 for the simulation under dilute coverage. That is, to estimate the preferred curvature of a patch at a protein coverage different from that simulated, the sampled curvature is multiplied by the increase in protein coverage from the simulated system to the target. Note here that 𝐴_p_ cancels when translating from 〈𝐽_protein_〉_sim_ to 𝐽_0,target_.

With this method, spontaneous curvature is best estimated from simulations where the protein has low coverage and is relatively small compared to the undulation wavelength. At complete coverage, the protein will cover the surface homogeneously and would be unable to enrich at the curvature it induces, yielding little information. We therefore conducted low coverage simulations for dimeric, tetrameric, and hexameric peripherin-2 linear polymer chains in a POPC bilayer model. Cartoon representations of MD simulations for a non-covalent dimer (Fig. 7D) and for a hexamer chain (Fig. 7E) species show representative views. Figure 7F plots the curvature sampled, per mode, by each of the three species (dimer, tetramer chain, hexamer chain) simulated. Here, the curvature sampled indicates the dilute simulation patch spontaneous curvature as 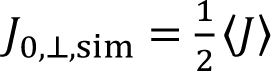. Disk rim patch spontaneous curvature is modeled to be

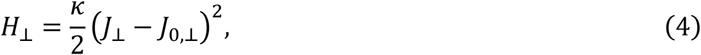

where spontaneous curvature is extrapolated from the dilute simulation coverage (ϕ_sim_ = 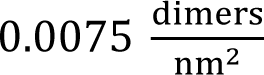 for the hexamer) to the actual disk rim coverage 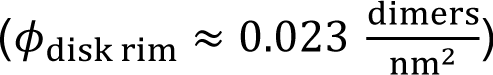, estimated using a peripherin-2 dimer repeat distance of 4.1nm from (28).

The MD simulation findings (Fig. 7F) show that curvature sampled varies roughly linearly with the number of dimers, and that the tetramer and hexamer chain forms each generate anisotropies far greater than those produced by individual dimers. For all forms except the loosely restrained dimer, strong negative curvature was induced perpendicular to the chain axis, while only weak curvature was induced in the parallel direction. In the biological context, this predicts the generation of extreme disk rim membrane bending towards the peripherin-2 EC2 domains and away from the OS cytoplasm. The spontaneous rim curvature implied by the MD simulations of the tightly restrained model is ca. 0.079 nm^-1^, which is equivalent to a disk rim diameter of ∼25 nm measured at the bilayer midplane (ca. 27 nm measured at the OD). This is notably close to the 24 nm OD of disk rims observed experimentally, and suggests that extreme membrane curvature can be generated by a high coverage of individual (non-associating) peripherin-2 belts. For the loosely restrained model, curvature induction is roughly halved (Fig. 7F). These findings suggest that protein shape rigidity is an important parameter for membrane curvature generation by peripherin-2.

## Discussion

This report advances knowledge and understanding of peripherin-2 and rom1, photoreceptor-specific tetraspanins that assemble into a supramolecular transmembrane scaffold that sculpts the distinctive rim domains of rod and cone OS disks. Because the detailed organization of the disk rim scaffold - and the mechanism by which it generates extreme membrane curvature - is not yet known, we applied velocity sedimentation, BN-PAGE, and computational molecular modeling approaches to address these gaps. Our findings suggest that normal assembly of the peripherin-2/rom1 disk rim scaffold includes at least four types of protein-protein interactions that together support normal OS structure. Moreover, they reveal that peripherin-2 non-covalent dimers that are incorporated into disulfide-linked chains generate highly anisotropic negative membrane curvature – the defining geometric feature of disk rim structure. These considerations suggest that rim formation during disk morphogenesis includes the incorporation of peripherin-2 chains and free dimers into three parallel and interacting belts that gradually lengthen to propel disk rim advance and drive disk internalization towards completion. Importantly, this work also establishes new experimental and computational approaches to systematically evaluate and rank the pathogenicity of *PRPH2* mutations, thereby enabling future mechanistic studies of associated IRDs.

Investigations of peripherin-2 and rom1 have relied heavily on velocity sedimentation to characterize detergent-solubilized forms of the proteins (2). Originally developed to assess the fidelity of recombinant expression and the impact of mutations on peripherin-2 and rom1 protein structure (18,34,36,38), the technique has since been the method of choice to investigate peripherin-2 and rom1 self-assembly in numerous engineered mouse models (2,42). Although this approach can reveal gross changes in protein “complex formation”, limitations for its tractability, reproducibility, and dynamic range motivated us to redesign it. Because the first two shortcomings are a function of a challenging manual fractionation of small-volume (2 ml) sucrose density gradients, we developed a semi-automated technique to surmount them and reduced g-force to improve assay dynamic range. The new experimental design reduced the level of manual dexterity required for gradient collection, improved reproducibility, and prevented signal clipping caused by pelleting, thereby improving assay dynamic range.

The reoptimized method generated sedimentation profiles that highlight the broad, stable, and continuous distribution of peripherin-2 and rom1 polymeric forms that are released from ROS membranes by gentle detergent solubilization. To advance understanding of the polymers observed in sedimentation experiments, we developed a BN-PAGE strategy that reports on the absolute chain length and relative abundance of peripherin-2 dimer chains. The BN-PAGE assay identified a range of peripherin-2 and rom1 dimer chain lengths in bovine and murine ROS membranes that are in good agreement with each other, and with previous findings from negative-staining TEM and single-particle cryo-EM studies (21,29). Application of BN-PAGE to sedimentation gradient fractions emphasized that, while sedimentation effectively distributes heterogeneously-sized polymers throughout sucrose density gradients, it fails to fully resolve different dimer chain species.

The availability of an efficient BN-PAGE assay for peripherin-2/rom1 polymer assembly has significant clinical implications. Three decades have elapsed since velocity sedimentation was first used to identify a protein-level defect caused by a pathogenic mutation in *PRPH2* (41), and more than 800 genetic variants have been identified in the interim (https://www.ncbi.nlm.nih.gov/clinvar). Clinical classifications, based on a framework developed by the American College of Medical Genetics and Genomics / Association for Molecular Pathology (ACMG/AMP), lack reliability and include a high proportion of variants of uncertain significance (VUS). A high-confidence, high-throughput classification method for *PRPH2*-associated disease mutations would therefore be of significant clinical benefit to aid patient counseling and the development of therapeutics (12). The findings reported here demonstrate the utility of BN-PAGE for peripherin-2 mutant screening. Work is in progress to scale this approach to catalog how *PRPH2* variants impact protein structure and function.

As a complement to recombinant protein studies, engineered mouse models play critical roles for understanding *PRPH2*-associated disease pathology and for the development of effective therapeutic strategies. Here, we used knockout mice together with velocity sedimentation and BN-PAGE to show that rom1 expression supports the assembly of the highest-order peripherin-2 polymers, a conclusion previously reached by Lewis et al (42). The mechanism by which rom1 promotes peripherin-2 polymerization is unclear, because rom1 is conspicuously absent from higher-order polymers (Figs. 1B, 2A-C). Lewis, et al suggest that rom1 may promote peripherin-2 polymerization by facilitating an optimal peripherin-2 internal disulfide bond arrangement (51). We show here that ROS membranes from *Rom1* KO mice include disulfide-linked peripherin-2 chains of up to 5 dimers, and we find that the proportion of peripherin-2 that is disulfide-bonded remains unchanged relative to that in WT mice (Fig. 6A). Thus, disulfide-mediated dimer polymerization is at least partially intact in the absence of rom1 and additional studies are needed to resolve this issue. Nonetheless, the results reported here demonstrate the utility of BN-PAGE as an effective method for molecular phenotyping of peripherin-2 in IRD model mice.

The biological significance of the longest peripherin-2 chains (comprising >5 dimers) remains unclear. In addition to the loss of these species in *Rom1* KO mouse photoreceptors, Lewis et al reported reduced disk internalization rates and an absence of disk incisures - despite a compensatory increase in peripherin-2 expression (42). Because the latter two phenotypes could be rescued by transgenic overexpression of peripherin-2, current evidence suggests that rom1 is redundant with peripherin-2 and its unique functional role, if any, remains to be determined.

It is well-established that peripherin-2 undergoes multistep self-assembly (20,21) in support of a supramolecular transmembrane rim scaffold that establishes and maintains OS disk rim curvature (28,29); however, the precise assembly pathway, and a resolved map of the protein-protein interactions that support the scaffold are still needed. Although early investigations proposed non-covalent tetramers as minimal structural elements (32,52), a single-particle cryo-EM study has since resolved non-covalent dimers as the minimal unit of peripherin-2 structure (29). That study illustrates that the non-covalent dimerization interface is formed by association of the two intradiskal EC2 ectodomains, which give the appearance of kissing heads. The two four-helix transmembrane bundles project through the membrane to its cytoplasmic surface, where C-termini project from the back of each monomer, giving tail-like appearances (see Fig. 5H; WT). This dimeric structure is consistent with structural evidence for other tetraspanins (53,54).

The studies reported here now suggest that non-covalent dimers can themselves spontaneously associate non-covalently in a side-by-side manner to form rows (Fig. 5A, 7A, B). These interactions appear to be of modest strength because they are disrupted by the gentle detergents used here. Because the dimer-dimer interface lies in a plane that includes C150 in peripherin-2 (and C153 in rom1), we hypothesize that the non-covalent organization of dimers into *rows* brings pairs of cysteines (contributed from different dimers) into a proximity that promotes the formation of disulfide-bonded dimer *chains*. The presence of dimer chains of varied lengths in mammalian ROS and transfected HEK293 cell membranes has been documented previously and is confirmed again here using an independent method (Fig. 3A, 5A). Overall, the new data suggest that the three parallel belts of transmembrane particles localized at disk rims (28) are comprised of a mixture of free peripherin-2 dimers and disulfide-linked dimer chains of varied lengths. Future studies will be needed to detail how peripherin-2/rom1 heterodimers are organized within these structures.

As a first step towards understanding how peripherin-2 supports the extreme curvature of disk rims, we conducted coarse-grained MD simulations using a strategy that infers spontaneous curvature from the dynamic position of the protein model on undulatory normal modes, where it occupies sites of high bilayer curvature. This approach, which interprets curvature on a per-normal-mode basis is straightforward to model and is insensitive to leaflet lipid-number asymmetry (50). It was therefore selected over simulation strategies applying static buckles (55–57), bicelles (58), and complex continuum-modeling (59). Our simulations predict that belts of peripherin-2 homodimer chains arrayed around the disk circumference generate substantial anisotropic negative membrane curvature, which curves away from the OS cytoplasm at an orientation perpendicular to the belt axis. This deformation is therefore predicted to bend the disk membrane to enclose peripherin-2 EC2 domains, while leaving N-and C-termini exposed in the cytoplasm – precisely the topology adopted in OSs. Importantly, when adjusted for the very high surface density of peripherin-2 at disk rims, MD simulations predict that *individual* belts containing peripherin-2 dimer chains generate significant membrane curvature. This finding has two interesting implications. First, it suggests that curvature generated by chains during disk rim morphogenesis may create energy wells that draw free dimers into alignment, amplifying the dimers’ weak tendency towards self-association. Second, because the energetic contributions of three individual (non-interacting) belts by itself may be adequate to produce extreme membrane curvature, interactions between the peripheral and central belts may play a relatively lesser role for scaffolding rim structure. Future computational modeling may help resolve this issue.

The coarse-grained Martini model used here is not designed to predict protein folds (often requiring atomic-level specificity), but rather to answer questions regarding collective behavior at long length-scales. We therefore compared loosely and tightly restrained models for the peripherin-2 homodimer. In each case (secondary or tertiary) sequence restraints facilitate protein model folding and stability, though to differing extents. The weaker curvature induction inferred with the loosely restrained model suggests that protein shape is critical for enforcing extreme membrane curvature. We expect that higher-order assembly of peripherin-2 into a supramolecular scaffold reinforces individual dimer shape by preventing conformations or fluctuations inconsistent with the triple-belted structure. However, current uncertainty regarding dimer shape rigidity is an important limitation of the present findings, and investigating protein dynamics will be an important goal for future work using all-atom simulations. In addition, the MD simulations reported here were performed using a bilayer of simplified composition (POPC) to facilitate simulations and minimize assumptions. Importantly, OS disks represent a distinctive membrane environment noted for high levels of phosphatidylethanolamine (PE) relative to the plasma membrane (60). Likewise, disk rims are reported to be enriched in PE relative to the OS plasma membrane (61). The conical PE headgroup tends to soften membranes and reduce the energy required to generate membrane curvature (62). Because POPC tends to resist curvature, results from the simplified approach used here are best viewed as a lower limit approximation for curvature generation induced by peripherin-2 *in situ*. These findings raise interesting new questions, and future work using all-atom simulations could address potential impacts of bulk lipid composition and of specific lipid-protein interactions. The observation that peripherin-2 expression in HEK293 induces extreme membrane curvature comparable to that of disk rims suggests that the distinctive lipid profile of ROS membranes is not required for its function (20,21).

In sum, this report advances our understanding of how multiple levels of peripherin-2 and rom1 protein-protein interactions assemble a supramolecular rim scaffold that sculpts the extreme curvature of photoreceptor OS disk rims. Like other superfamily members, peripherin-2 and rom1 engage in a web of protein-protein interactions that serve to organize a membrane microdomain (5,7). The findings, based on a combination of experimental and computational methods, provide a new model for how peripherin-2 stabilizes the extreme curvature required for photoreceptor OS disk structure. They also establish a potent molecular phenotyping strategy for peripherin-2 and rom1, which has substantial clinical significance, because mutations in *PRPH2* and *ROM1* are responsible for a broad variety of blinding IRDs. Because incomplete knowledge of *PRPH2* structure–function relationships currently hinders interpretation of genotype–phenotype relationships and classification of variants of uncertain significance (10,11,63), we anticipate that application of the new tools described here can help surmount these barriers.

## Experimental procedures

### Materials

Reagents and chemical were purchased from MilliporeSigma (St. Louis, MO) or Fisher Scientific (Waltham, MA) unless otherwise stated. Purified dark-adapted bovine ROS were obtained from InVision BioResources (Seattle, WA). Triton X-100 and 10:1 n-dodecyl-ß-D-maltoside (DDM) / Cholesteryl hemisuccinate (CHS) were from Anatrace (Maumee, Ohio). Mini-PROTEAN precast TGX gels and Coomassie Brilliant Blue (CBB) G-250 were purchased from BioRad Laboratories (Hercules, CA). HEK293 cells (a more adherent sub-line of ATCC CRL-1573 designated as AD293) were originally purchased from Stratagene (La Jolla, California) and were maintained in-house from frozen stocks. Codon-optimized plasmids for protein expression in HEK293 were generated by ATUM (Newark, California) and were sequence-verified prior to use.

### Animal care and use

Mouse care and use was in accordance with the ARVO Statement for the Use of Animals in Ophthalmic and Vision Research and was accredited by the Oakland University Institutional Animal Care and Use Committee. Mice were housed in a facility approved by the Association for the Assessment and Accreditation of Laboratory Animal Care International (AAALAC). Mice were maintained under 12h light/12h dark light cycles with food and water provided ad libitum; studies used roughly equal numbers of age-matched male and female mice. For tissue collection, mice were euthanized under normal room illumination by asphyxiation with CO_2_, followed by cervical dislocation.

The *Rom1* knockout mouse line (strain #4510) was obtained from The Jackson Laboratory (Bar Harbor, Maine) and was outcrossed to CD-1 mice (strain #022) from Charles River Laboratories (Wilmington, Massachusetts) for at least two generations to improve fecundity. The line was confirmed to be free of common IRD mutations, by genotyping at the *rd1* (Pde6b) and *rd8* (Crb1) loci. Mice carrying the Rom1 mutation were identified via SYBR green qPCR melting curve assay using mix 1 (5’-TGCCCGTCTTTATCTTCCAG and 5’-CCTTCCACATTGCTCTGGAT) and/or mix 2 (5’-TTGGGTGGAGAGGCTATTCG and 5’-CTTCCCGCTTCAGTGACAAC) primers. SDS-PAGE was used to confirm a complete absence of rom1 in ROS prepared from rom1 KO retinas.

### Cell culture and recombinant protein expression

HEK293 cells were maintained in DMEM media containing 10% FBS and 5% penicillin/streptomycin, using under standard culture conditions of 37^0^C with 5% CO_2_. Transient transfections of HEK293 cells with X-tremeGENE 9 reagent (Roche, MilliporeSigma) were performed according to the manufacturer’s specifications and essentially as described previously (20,21), using cells subjected to fewer than 20 passages. Expression plasmids included a full-length peripherin-2 coding sequence, for WT bovine peripherin-2 or one of three variants - C150S, L185P, or C150S;L185P.

### Murine ROS preparation

Enucleations were performed immediately following mouse sacrifice. Retinas were isolated by dissection, flash-frozen in liquid nitrogen, and stored at -80°C until use. Pooled retinas (four per group) were thawed in 200 µl of chilled Resuspension Buffer (1x PBS; 136 mM NaCl, 2.7 mM KCl, 5.3 mM Na_2_HPO_4_, 0.7 mM, pH 7.4), supplemented with 50 mM N-ethylmaleimide, and 1x cOmplete ULTRA EDTA-free protease inhibitor cocktail. Retinas were vortexed to dislodge their OSs, centrifuged at room temperature (RT) for one minute at 500 rcf, and the supernatant was collected. The pelleted retinas were then resuspended and re-extracted two additional times with 200 µl of chilled Resuspension Buffer each time. The ROS-containing supernatants were pooled and pelleted by centrifugation at RT for two minutes at 10k rcf. The final ROS pellets were triturated in 50 µl of Resuspension Buffer, and the total protein concentration was estimated using A_280_ in Resuspension Buffer + 1% SDS.

### Protein solubilization

Bovine or murine ROS membranes were resuspended at 3 mg/ml in chilled Resuspension Buffer. An equal volume of chilled 1x PBS containing either 2% Triton X-100 or 2% DDM-CHS, was gradually added with gentle vortexing, to give final detergent and protein concentrations of 1% and 1.5-2 mg/ml respectively. Membranes were extracted on ice for one hour, then were centrifuged at 45k rpm (90k rcf) for 30 minutes at 4°C. Supernatants (“detergent extracts”) were collected and either used immediately, or stored at 4°C for use the following day. A similar process was followed for HEK293 cell membranes; however, cell pellets were resuspended at 20 mg/ml and solubilized at 10 mg/ml final total protein.

### Velocity sedimentation analyses

Velocity sedimentation of detergent extracted proteins from ROS or HEK293, in 5-20% (w/w) sucrose gradients, was performed using three modifications to the original method (18). First, DTT was omitted from all solutions (34). Second, sedimentations were performed at 25K rpm (41.5K rcf). Finally, gradient fractions were collected using a semi-automated method, employing custom-designed tools for tube puncture and capping. Detailed methods are described in the Supporting Information and Figure S1. Fractionated gradients were analyzed by non-reducing SDS-PAGE as described previously (20), and/or by BN-PAGE (described below) and western blotting.

Western blotting was performed essentially as described (64). Primary antibodies included anti-mouse peripherin-2, MabC6 (30) and PabMPCT (65); anti-bovine peripherin-2, PabBCT (66); and the anti-rom1 antibodies Mab2H5 (Millipore MABN1757) and PabMUTT (30). Fluorescent secondary antibodies included goat anti-mouse and goat anti-rabbit IRDye 680RD and IRDye 800CW (Li-Cor Biosciences, Lincoln, NE), and goat anti-rabbit AS490 and AS550 (Azure Biosystems, Dublin, CA). Immunoblots were scanned using a Sapphire Biomolecular Imager (Azure Biosystems, Dublin, CA) and densitometry was performed using Image J or AzureSpot software. Cumulative sedimentation profiles (Fig. 4C, D) were modeled using locally estimated scatterplot smoothing (LOESS) regression in RStudio. Optimal LOESS spans for each experimental group were selected by minimizing the Akaike Information Criterion (AIC).

### BN-PAGE electrophoresis and transfer

Blue native PAGE (BN-PAGE) (67,68) was optimized for resolving peripherin-2 and rom1-containing polymers from ROS and HEK293 cell membranes. Soluble detergent extracts from either ROS or HEK293 were combined 2:1 with 3x BN-PAGE sample buffer (100 mM Tris, 60% glycerol, 0.15% bromophenol blue, 0.25% CBB G-250, 25 mM 6-animohexanoic acid (ε-Ahx), pH 7.2) and loaded onto 4-20% TGX gradient gels. Electrophoresis was performed on ice using chilled Cathode buffer B (Tris-Glycine running buffer + 0.02% CBB G-250 and 2 mM ε-AHX) and chilled Anode buffer (25 mM Tris + 192 mM glycine, pH 8.3). Gels were run at 100V for 30 minutes, and then at 150V until the dark blue dye front had migrated 1/3 of the way into the gel. Electrophoresis was paused and Cathode buffer B was replaced with buffer B/10 (Tris-Glycine running buffer + 0.002% CBB G-250 and 0.2 mM ε-Ahx). Electrophoresis was resumed at 100V until the dark blue dye front reached the bottom of the gels. Following electrophoresis, gels were equilibrated at RT in Anode buffer containing 2% SDS for 30 minutes, and then in Anode buffer alone for 3x 5 minutes. Separated proteins were transferred to Immobilon-FL membranes as previously described (20), and membranes were rinsed in ultrapure water and air dried. Prior to development, membranes were rinsed in 100% methanol, equilibrated 15 minutes in 40% methanol + 7% acetic acid, rinsed in ultrapure water, and equilibrated in 1xPBS. Standard immunoblotting methods (described above for SDS-PAGE western blots) were used to detect proteins on membranes. Species apparent molecular weights were calculated from logarithmic regression of NativeMark protein standards.

### Molecular dynamics simulations

Coarse-grained (CG) MD simulations of peripherin-2 dimers, tetramers, and hexamers embedded in flat membrane patches were prepared from AlphaFold2 multimer all-atom structures using the Martini Maker tool from CHARMM-GUI (69). Martini 2 was used as the CG model (46,70). The bilayer lipids consisted of 1200 POPC lipids (with 8 more lipids in the upper leaflet for each dimer). A pure POPC bilayer model was selected for MDS approximation of protein curvature induction because it avoids the extensive simulation times required for a mixture of curvature-sensitive lipids to relax their dynamic distribution on a slow diffusion timescale. Because curvature-favoring lipids are absent, this simplified approach is expected to provide a lower limit estimate of curvature-generation induced by protein shape in a relatively curvature-resistant bilayer model. Additionally, because this model lacks other lipid species and uses a coarse-grained force field, it does not address the potential for particular peripherin-2-interacting lipids to influence curvature induction.

The simulations were run using GROMACS (71) with the default Martini 2 parameters: a timestep of 20 fs, 1.1 nm cutoff for non-bonded interactions, v-rescale temperature coupling at 310.15 K, and semi-isotropic Parrinello-Rahman pressure coupling with 12 ps time constant and compressibility of 3×10^-4^ bar^-1^. The C150 disulfide bonds were included between dimers using harmonic restraints of length 0.39 nm and strength 5000 kJ/mol/nm^2^, and the proteins were aligned along the Y axis with a harmonic restraint along the X axis between the first and last dimers in the tetramer and hexamer simulations, and a restraint along Y between the molecules in the dimer simulation. Each unconstrained system was executed 10 μs for each of 5 replicas. To keep the overly flexible (loosely restrained) Martini coarse-grained model consistent with the AlphaFold starting structure, a tightly restrained model was also studied, in which root-mean-squared deviation (RMSD) restraints were applied with the Plumed software plugin. The force constant on the RMSD from the AF2 structure was 1000 kJ/mol/nm^2^, allowing a typical fluctuation of the RMSD of about 0.7 Angstroms. The performance penalty with the external Plumed software package necessitated more replicas that were run for shorter duration (20 replicas, 2.5 ms per replica).

### Spontaneous curvature analysis

Curvature was analyzed with the MembraneAnalysis Julia package (72). For each frame, Fourier mode amplitudes are analyzed using the approximate neutral surface atom of the lipids. The center-of-mass of the protein complex is used to evaluate its lateral position. Curvature sampled is then evaluated separately for perpendicular and parallel membrane undulations, using the two highest wavelength modes.

Eq. 1 is invariant to the local coordinate system, as 𝐽 = 𝐽_..l_ + 𝐽_∥_ and 𝐾 = 𝐽_..l_𝐽_∥_ are invariant. In the case of a large protein complex like peripherin-2, rotational symmetry is broken and spontaneous curvature is anisotropic. That is, spontaneous curvature varies either perpendicular or parallel (as indicated by 𝐽_0,..l_ and 𝐽_0,∥_, respectively) to the dimer, tetramer, or hexamer. Curvature perpendicular and parallel to the polymer axis is characterized separately and modeled by independent, *oriented* versions of Eq. 1,

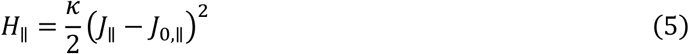

and

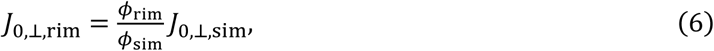

## Data availability

Data are contained within the manuscript and Supporting Information, with the exception of the MD simulation dataset, which can be accessed at https://doi.org/10.5281/zenodo.19049042.

## Acknowledgements

The authors express their utmost appreciation to Dr. Michael Latcha and his Senior Design Team (Oakland University Department of Mechanical Engineering) for development and fabrication of the semi-automated gradient fractionation apparatus and to Heather McDonald for illustrations.

This work was supported by the National Institutes of Health grant EY025291 and the Foundation Fighting Blindness award TA-GT-0424-0883-OAK to AFXG. AH, AHB, and AJS were supported by the Intramural Research Program of the *Eunice Kennedy Shriver* National Institute of Child Health and Human Development, National Institutes of Health (ZIA-HD008955). Molecular dynamics simulations were performed using the computational resources of the NIH High-Performance Computing Biowulf cluster (https://hpc.nih.gov).

## Abbreviations

DTT: dithiothreitol
DDM: docecylmaltoside
EC2: extracellular 2
LOESS: locally estimated scatterplot smoothing
OS: outer segment
ROS: rod outer segment

## SUPPORTING INFORMATION (Boucher et al)

**Figure S1.**
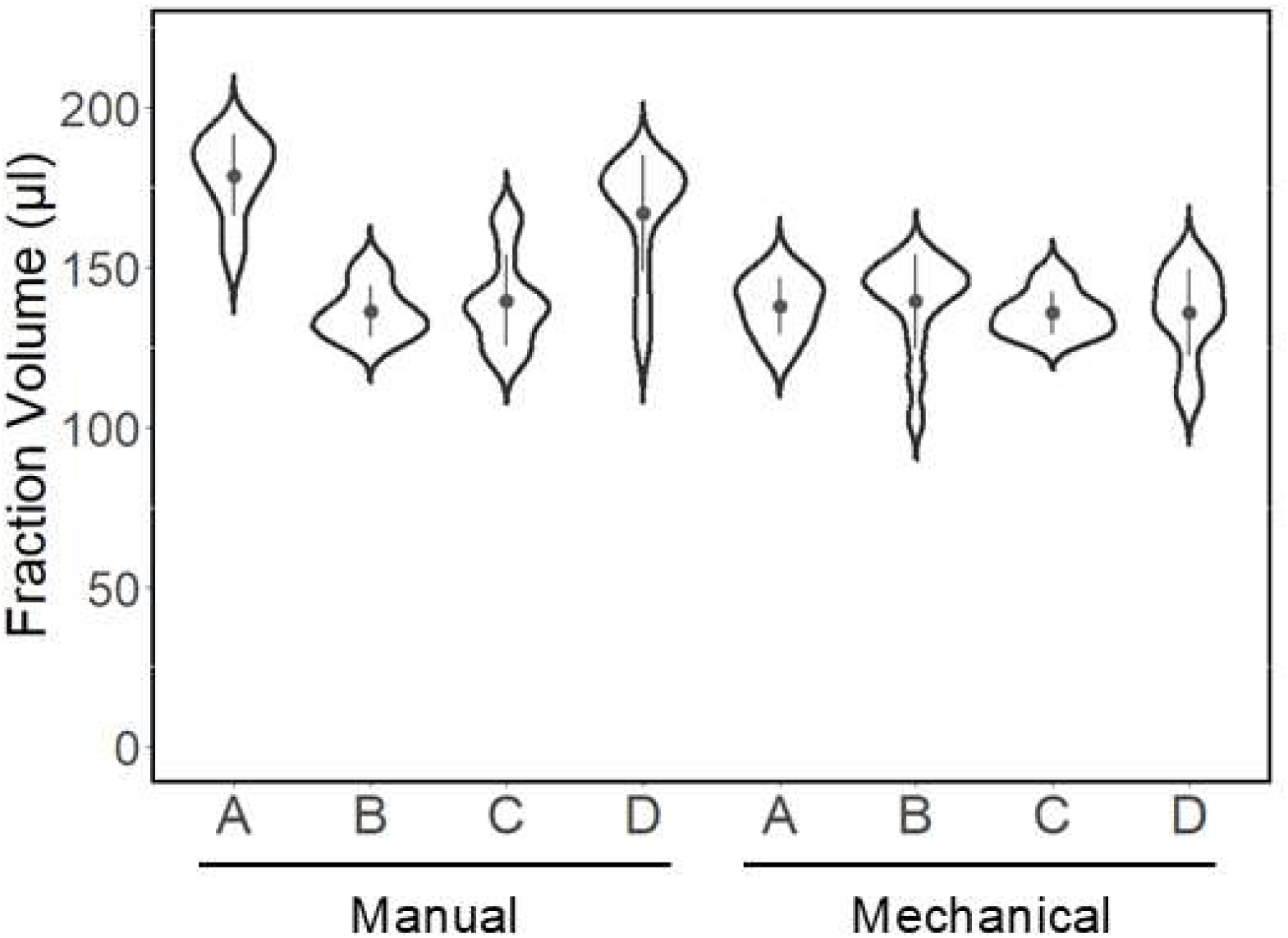
Mechanical fractionation reduces fraction volume variability. Violin plot comparing fraction volumes from four manually fractionated and four mechanically fractionated Triton X-100 sucrose gradients. Fraction volumes were measured using a P200 micropipette immediately following collection. The number of fractions collected varied based on the method used. Manually fractionated gradient produced as few as 11 and as many as 15 fractions, whereas mechanically fractionated gradients generally produced 14 to 15 fractions. Mechanical fractionation produced a slight linear increase in fraction volume (1.67*10^-2^ µl increase per µl collected, or 10.9 µl increase per cm from the bottom of the gradient, corresponding to ∼2.2 µl increase per successive fraction; R^2^ = 0.72, P < 10^-4^). Manual fractionation showed much greater variability in fraction volume, with no statistically significant correlation between fraction volume and position in the gradient. Normality of fraction volumes was assessed by Shapiro-Wilke test (P = 9.2*10^-3^ for mechanical fractionation, P < 10^-4^ for manual), indicating non-normal distributions. Variance in fraction volume was reduced using mechanical vs. manual fractionation (Fligner-Killeen test, P < 10^-4^).

**Figure S2.**
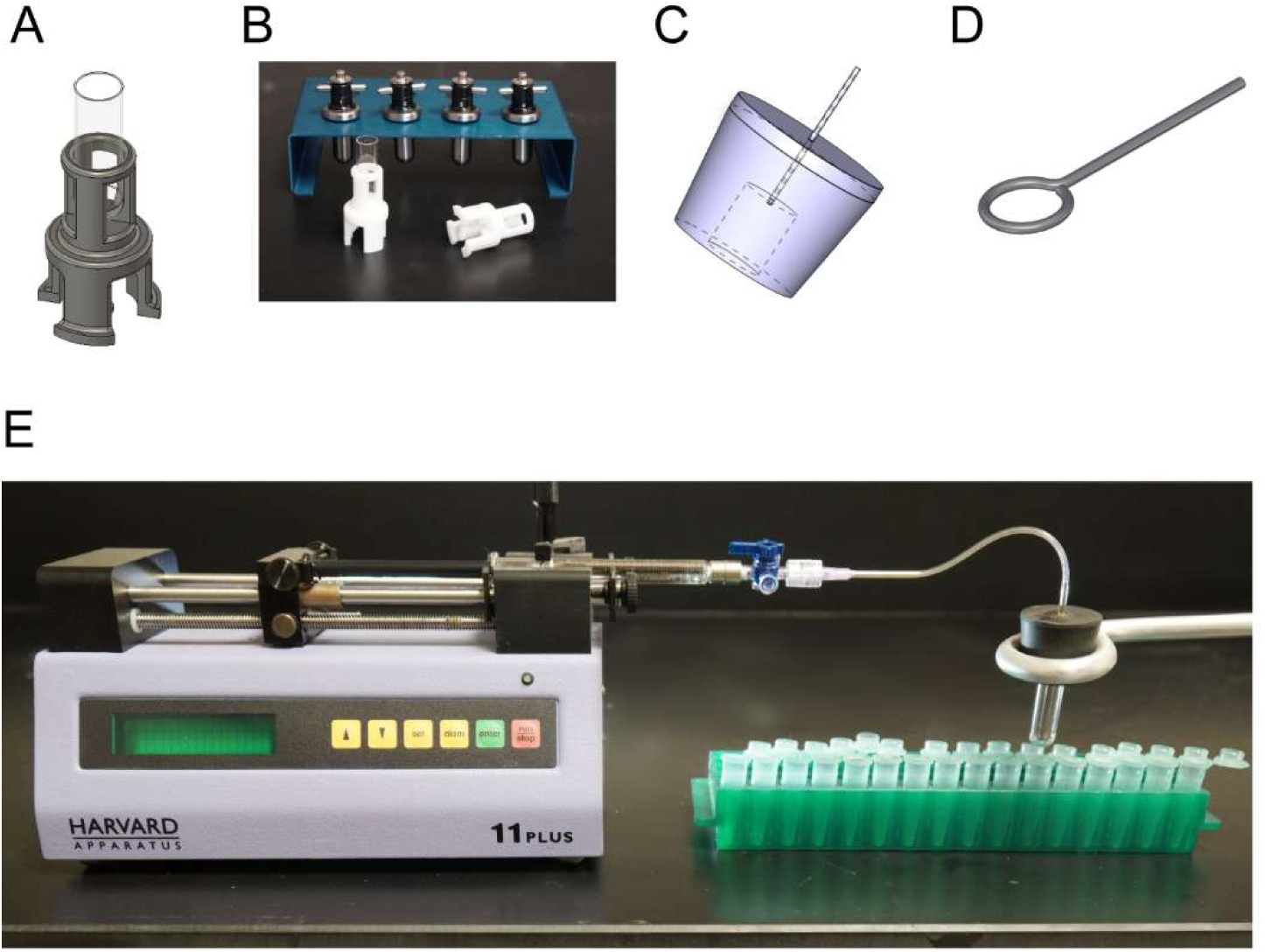
Apparatus for mechanical fractionation of 2 ml sucrose gradients. A) Final design of a 3D-printable tube punch, used to reproducibly pierce the bottom of 2.2 ml Open-top Thinwall Ultra-Clear (11×34mm) centrifuge tubes (Beckman, Brea, CA) used for sedimentations. B) Examples of 3D-printed tube punches, showing one (*right*) with a 27G1/2 syringe needle installed. A 3D (STL) print file for the tube punch is provided as a separate SI file. C) Design for the neoprene tube sealing cap, bored to snugly accommodate an inserted centrifuge tube. D) Design for a ring stand-mountable sealing cap support. E) Overview showing the precision syringe pump used to regulate mechanical fractionation connected to the sealing cap via a 3-way luer-lock airtight valve. Detailed procedures for use are given below.

**Figure S3.**
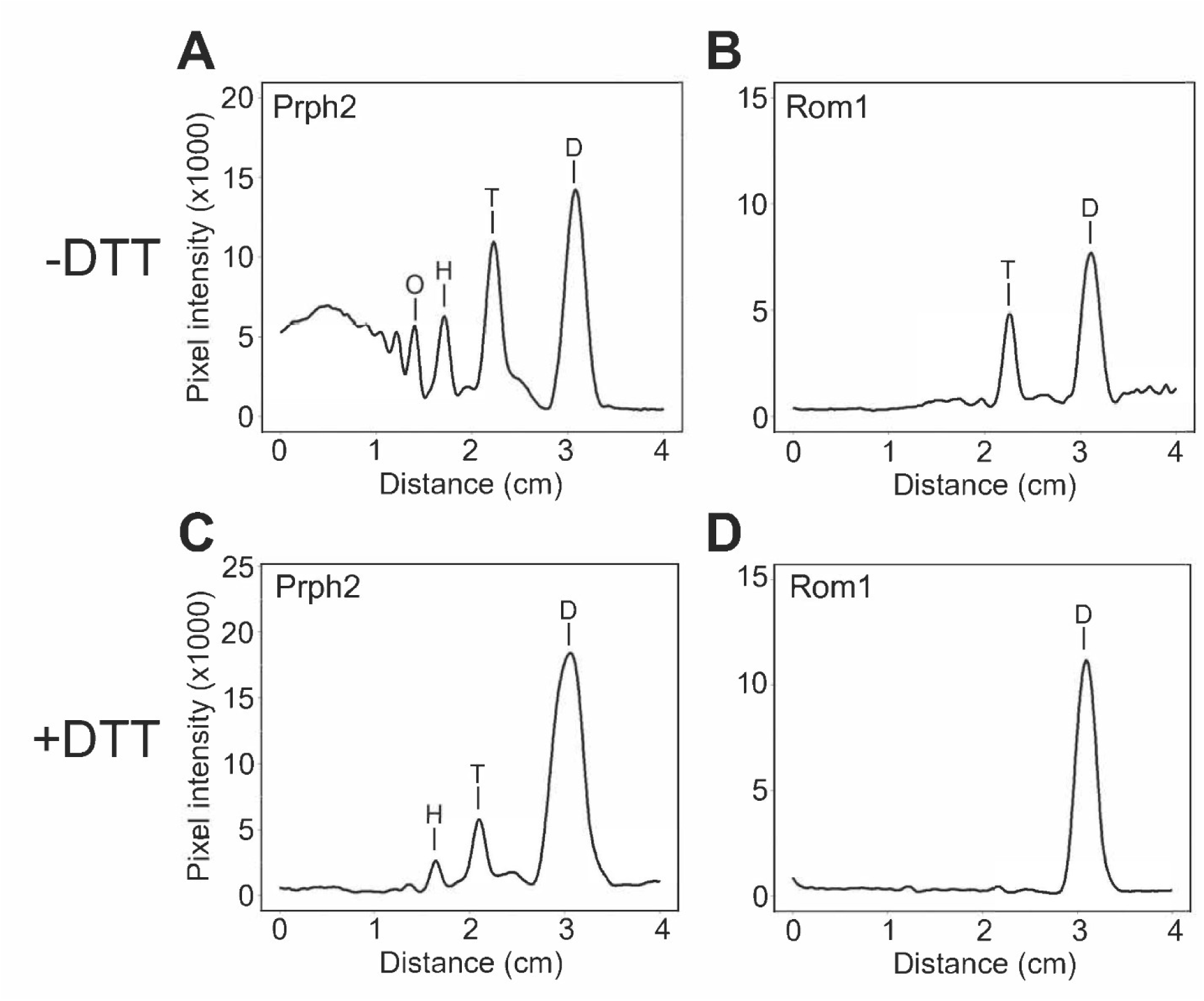
BN-PAGE assay reveals relative abundance of Prph2 and Rom1 polymers solubilized from bovine ROS membranes with Triton X-100. (A, B) Lane scan profiles of western blot bands for native Prph2 and Rom1. (C, D) Analogous data is shown for Prph2 and Rom1 reduced with DTT prior to BN-PAGE analysis. D, dimer; T, tetramer; H, hexamer; O, octamer.

## Detailed procedures for mechanical sucrose gradient fractionation

1. Following centrifugation, TLS-55 buckets were carefully transferred to their storage rack on ice. Gradients remained on ice until immediately prior to fractionation and were handled exclusively with forceps to minimize warming.
2. The tube punch was prepared with a new 27G1/2 syringe needle sealed with parafilm.
3. The syringe pump was prepared with a 2.5 ml luer-lock Gastight Hamilton syringe, a 3-way luer-lock valve, Tygon tubing, and a sealing cap. A rack was prepared with fifteen 1.5ml microcentrifuge collection tubes.
4. A thin layer of vacuum grease was applied to the bore of the sealing cap.
5. A gradient was removed from its bucket and the air-vented sealing cap was carefully pressed onto it.
6. The capped tube was carefully transferred to the punch, and gently but firmly pressed down to drive the needle through its bottom.
7. The 3-way valve was then closed to isolate the sealing cap and the capped tube was transferred to the ring stand support. The first collection tube was positioned directly underneath the punched hole.
8. The valve was switched to connect the syringe to the sealing cap and the pump was started at a constant flow rate of 1 ml/min, and 6 drop (∼140 µl) fractions were collected. The final fraction was often partial volume, which was noted.
9. When the gradient was fully expelled, fraction volume measured with a P200 micropipette, then tubes were sealed and stored at 4°C until use.

## References

1. Goldberg, A. F., Moritz, O. L., and Williams, D. S. (2016) Molecular basis for photoreceptor outer segment architecture. Prog. Retin. Eye Res. 55, 52–81. 10.1016/j.preteyeres.2016.05.003

2. Tebbe, L., Kakakhel, M., Makia, M. S., Al-Ubaidi, M. R., and Naash, M. I. (2020) The Interplay between Peripherin 2 Complex Formation and Degenerative Retinal Diseases. Cells. 10.3390/cells9030784

3. Dowling, J. E. (1987) The retina: an approachable part of the brain, Harvard University Press, Cambridge, MA

4. Hemler, M. E. (2005) Tetraspanin functions and associated microdomains. Nature Reviews Molecular Cell Biology 6, 801–811. DOI 10.1038/nrm1736

5. Charrin, S., Jouannet, S., Boucheix, C., and Rubinstein, E. (2014) Tetraspanins at a glance. J. Cell Sci. 127, 3641–3648. 10.1242/jcs.154906

6. Charrin, S., le, N. F., Silvie, O., Milhiet, P. E., Boucheix, C., and Rubinstein, E. (2009) Lateral organization of membrane proteins: tetraspanins spin their web. Biochem. J. 420, 133–154. BJ20082422 [pii];10.1042/BJ20082422 [doi]

7. van Deventer, S. J., Dunlock, V. E., and van Spriel, A. B. (2017) Molecular interactions shaping the tetraspanin web. Biochem. Soc. Trans. 45, 741–750. 10.1042/BST20160284

8. Conley, S. M., Stuck, M. W., and Naash, M. I. (2012) Structural and functional relationships between photoreceptor tetraspanins and other superfamily members. Cell Mol. Life Sci. 69, 1035–1047. 10.1007/s00018-011-0736-0 [doi]

9. Goldberg, A. F. X. (2013) Essential tetraspanin functions in the vertebrate retina. in Tetraspanins (Berditchevski, F., and Rubinstein, E. eds.), Springer Science+Business Media, Dordretcht. pp 321–344

10. Reeves, M. J., Goetz, K. E., Guan, B., Ullah, E., Blain, D., Zein, W. M., Tumminia, S. J., and Hufnagel, R. B. (2020) Genotype-phenotype associations in a large PRPH2-related retinopathy cohort. Hum Mutat 41, 1528–1539. 10.1002/humu.24065

11. Peeters, M., Khan, M., Rooijakkers, A., Mulders, T., Haer-Wigman, L., Boon, C. J. F., Klaver, C. C. W., van den Born, L. I., Hoyng, C. B., Cremers, F. P. M., den Hollander, A. I., Dhaenens, C. M., and Collin, R. W. J. (2021) PRPH2 mutation update: In silico assessment of 245 reported and 7 novel variants in patients with retinal disease. Hum Mutat 42, 1521–1547. 10.1002/humu.24275

12. Ayyagari, R., Borooah, S., Durham, T., Gelfman, C., and Bowman, A. (2024) Current and Future Directions in Developing Effective Treatments for PRPH2-Associated Retinal Diseases: A Workshop Report. Transl Vis Sci Technol 13, 16. 10.1167/tvst.13.10.16

13. Sanyal, S., and Jansen, H. G. (1981) Absence of receptor outer segments in the retina of *rds* mutant mice. Neurosci. Lett. 21, 23–26

14. Hawkins, R. K., Jansen, H. G., and Sanyal, S. (1985) Development and degeneration of retina in *rds* mutant mice: photoreceptor abnormalities in the heterozygotes. Exp. Eye Res. 41, 701–720

15. Clarke, G., Goldberg, A. F., Vidgen, D., Collins, L., Ploder, L., Schwarz, L., Molday, L. L., Rossant, J., Szel, A., Molday, R. S., Birch, D. G., and McInnes, R. R. (2000) Rom-1 is required for rod photoreceptor viability and the regulation of disk morphogenesis. Nat. Genet. 25, 67–73

16. Ma, C. J., Lee, W., Stong, N., Zernant, J., Chang, S., Goldstein, D., Nagasaki, T., and Allikmets, R. (2019) Late-onset pattern macular dystrophy mimicking ABCA4 and PRPH2 disease is caused by a homozygous frameshift mutation in ROM1. Cold Spring Harb Mol Case Stud. 10.1101/mcs.a003624

17. Bascom, R. A., Manara, S., Collins, L., Molday, R. S., Kalnins, V. I., and McInnes, R. R. (1992) Cloning of the cDNA for a novel photoreceptor membrane protein (rom-1) identifies a disk rim protein family implicated in human retinopathies. Neuron 8, 1171–1184

18. Goldberg, A. F., Moritz, O. L., and Molday, R. S. (1995) Heterologous expression of photoreceptor peripherin/rds and Rom-1 in COS-1 cells: assembly, interactions, and localization of multisubunit complexes. Biochemistry 34, 14213–14219

19. Arikawa, K., Molday, L. L., Molday, R. S., and Williams, D. S. (1992) Localization of peripherin/rds in the disk membranes of cone and rod photoreceptors: relationship to disk membrane morphogenesis and retinal degeneration. J. Cell Biol. 116, 659–667

20. Milstein, M. L., Kimler, V. A., Ghatak, C., Ladokhin, A. S., and Goldberg, A. F. X. (2017) An inducible amphipathic helix within the intrinsically disordered C terminus can participate in membrane curvature generation by peripherin-2/rds. J Biol Chem 292, 7850–7865. 10.1074/jbc.M116.768143

21. Milstein, M. L., Cavanaugh, B. L., Roussey, N. M., Volland, S., Williams, D. S., and Goldberg, A. F. X. (2020) Multistep peripherin-2/rds self-assembly drives membrane curvature for outer segment disk architecture and photoreceptor viability. Proc Natl Acad Sci U S A 117, 4400–4410. 10.1073/pnas.1912513117

22. Molday, R. S., and Moritz, O. L. (2015) Photoreceptors at a glance. J. Cell Sci. 128, 4039–4045. 10.1242/jcs.175687

23. Arshavsky, V. Y., and Burns, M. E. (2012) Photoreceptor signaling: supporting vision across a wide range of light intensities. J. Biol. Chem. 287, 1620–1626. R111.305243 [pii];10.1074/jbc.R111.305243 [doi]

24. Palczewski, K. (2012) Chemistry and biology of vision. J. Biol. Chem. 287, 1612–1619. R111.301150 [pii];10.1074/jbc.R111.301150 [doi]

25. Steinberg, R. H., Fisher, S. K., and Anderson, D. H. (1980) Disc morphogenesis in vertebrate photoreceptors. J. Comp. Neurol. 190, 501–508

26. Corless, J. M., Fetter, R. D., Zampighi, O. B., Costello, M. J., and Wall-Buford, D. L. (1987) Structural features of the terminal loop region of frog retinal rod outer segment disk membranes: II. Organization of the terminal loop complex. J. Comp. Neurol. 257, 9–23

27. Jarsch, I. K., Daste, F., and Gallop, J. L. (2016) Membrane curvature in cell biology: An integration of molecular mechanisms. J Cell Biol 214, 375–387. 10.1083/jcb.201604003

28. Poge, M., Mahamid, J., Imanishi, S. S., Plitzko, J. M., Palczewski, K., and Baumeister, W. (2021) Determinants shaping the nanoscale architecture of the mouse rod outer segment. Elife. 10.7554/eLife.72817

29. El Mazouni, D., and Gros, P. (2022) Cryo-EM structures of peripherin-2 and ROM1 suggest multiple roles in photoreceptor membrane morphogenesis. Sci Adv 8, eadd3677. 10.1126/sciadv.add3677

30. Goldberg, A. F., Fales, L. M., Hurley, J. B., and Khattree, N. (2001) Folding and subunit assembly of photoreceptor peripherin/rds is mediated by determinants within the extracellular/intradiskal EC2 domain: implications for heterogeneous molecular pathologies. J. Biol. Chem. 276, 42700–42706

31. Ding, X. Q., Stricker, H. M., and Naash, M. I. (2005) Role of the second intradiscal loop of peripherin/rds in homo and hetero associations. Biochem. 44, 4897–4904

32. Goldberg, A. F., and Molday, R. S. (1996) Subunit composition of the peripherin/rds-rom-1 disk rim complex from rod photoreceptors: hydrodynamic evidence for a tetrameric quaternary structure. Biochemistry 35, 6144–6149

33. Goldberg, A. F., Loewen, C. J., and Molday, R. S. (1998) Cysteine residues of photoreceptor peripherin/rds: role in subunit assembly and autosomal dominant *retinitis pigmentosa*. Biochem. 37, 680–685

34. Loewen, C. J., and Molday, R. S. (2000) Disulfide-mediated oligomerization of Peripherin/Rds and Rom-1 in photoreceptor disk membranes. Implications for photoreceptor outer segment morphogenesis and degeneration. J. Biol. Chem. 275, 5370–5378

35. Molday, R. S., Hicks, D., and Molday, L. (1987) Peripherin. A rim-specific membrane protein of rod outer segment discs. Invest. Ophthalmol. Vis. Sci. 28, 50–61

36. Loewen, C. J., Moritz, O. L., and Molday, R. S. (2001) Molecular characterization of peripherin-2 and rom-1 mutants responsible for digenic *retinitis pigmentosa*. J. Biol. Chem. 276, 22388–22396

37. Wittig, I., Beckhaus, T., Wumaier, Z., Karas, M., and Schägger, H. (2010) Mass Estimation of Native Proteins by Blue Native Electrophoresis PRINCIPLES AND PRACTICAL HINTS. Molecular & Cellular Proteomics 9, 2149–2161. DOI 10.1074/mcp.M900526-MCP200

38. Goldberg, A. F., and Molday, R. S. (2000) Expression and characterization of peripherin/rds-rom-1 complexes and mutants implicated in retinal degenerative diseases. Meth. Enzymol. 316, 671–687

39. Kajiwara, K., Berson, E. L., and Dryja, T. P. (1994) Digenic *retinitis pigmentosa* due to mutations at the unlinked peripherin/RDS and ROM1 loci. Science 264, 1604–1608

40. Seddon, J. M., De, D., Grunenkovaite, L., and Ferrara, D. (2024) Clinical and Imaging Characteristics of PRPH2 Retinopathies in a Longitudinal Cohort and Diagnostic Implications. Invest Ophthalmol Vis Sci 65, 31. 10.1167/iovs.65.14.31

41. Goldberg, A. F., and Molday, R. S. (1996) Defective subunit assembly underlies a digenic form of *retinitis pigmentosa* linked to mutations in peripherin/rds and rom-1. Proc. Natl. Acad. Sci. USA 93, 13726–13730

42. Lewis, T. R., Makia, M. S., Castillo, C. M., Hao, Y., Al-Ubaidi, M. R., Skiba, N. P., Conley, S. M., Arshavsky, V. Y., and Naash, M. I. (2023) ROM1 is redundant to PRPH2 as a molecular building block of photoreceptor disc rims. Elife. 10.7554/eLife.89444

43. Helfrich, W. (1973) Elastic properties of lipid bilayers: theory and possible experiments. Z Naturforsch C 28, 693–703. 10.1515/znc-1973-11-1209

44. Canham, P. B. (1970) The minimum energy of bending as a possible explanation of the biconcave shape of the human red blood cell. J. Theor. Biol. 26, 61–81. 10.1016/s0022-5193(70)80032-7

45. Steinkühler, J., Sezgin, E., Urbancic, I., Eggeling, C., and Dimova, R. (2019) Mechanical properties of plasma membrane vesicles correlate with lipid order, viscosity and cell density. Commun Biol 2. ARTN 337 10.1038/s42003-019-0583-3

46. Marrink, S. J., Risselada, H. J., Yefimov, S., Tieleman, D. P., and de Vries, A. H. (2007) The MARTINI force field: Coarse grained model for biomolecular simulations. Journal of Physical Chemistry B 111, 7812–7824. 10.1021/jp071097f

47. Mccammon, J. A., Gelin, B. R., and Karplus, M. (1977) Dynamics of Folded Proteins. Nature 267, 585–590. DOI 10.1038/267585a0

48. Monticelli, L., Kandasamy, S. K., Periole, X., Larson, R. G., Tieleman, D. P., and Marrink, S. J. (2008) The MARTINI coarse-grained force field: Extension to proteins. Journal of Chemical Theory and Computation 4, 819–834. 10.1021/ct700324x

49. Kroon, P. C., Grunewald, F., Barnoud, J., van Tilburg, M., Brasnett, C., Souza, P. C. T., Wassenaar, T. A., and Marrink, S. J. (2025) Martinize2 and Vermouth provide a unified framework for molecular topology generation. Elife 12. 10.7554/eLife.90627

50. Sapp, K. C., Beaven, A. H., and Sodt, A. J. (2021) Spatial extent of a single lipid’s influence on bilayer mechanics. Phys Rev E 103, 042413. 10.1103/PhysRevE.103.042413

51. Lewis, T. R., Phan, S., Castillo, C. M., Kim, K. Y., Coppenrath, K., Thomas, W., Hao, Y., Skiba, N. P., Horb, M. E., Ellisman, M. H., and Arshavsky, V. Y. (2023) Photoreceptor disc incisures form as an adaptive mechanism ensuring the completion of disc enclosure. Elife. 10.7554/eLife.89160

52. Kevany, B. M., Tsybovsky, Y., Campuzano, I. D., Schnier, P. D., Engel, A., and Palczewski, K. (2013) Structural and functional analysis of the native peripherin-ROM1 complex isolated from photoreceptor cells. J. Biol. Chem. 288, 36272–36284. M113.520700 [pii];10.1074/jbc.M113.520700 [doi]

53. Kovalenko, O. V., Yang, X. W., Kolesnikova, T. V., and Hemler, M. E. (2004) Evidence for specific tetraspanin homodimers: inhibition of palmitoylation makes cysteine residues available for cross-linking. Biochemical Journal 377, 407–417

54. Rubinstein, E., Théry, C., and Zimmermann, P. (2025) Tetraspanins affect membrane structures and the trafficking of molecular partners: what impact on extracellular vesicles? Biochem. Soc. Trans. 53, 371–382. 10.1042/Bst20240523

55. Mandal, T., Spagnolie, S. E., Audhya, A., and Cui, Q. (2021) Protein-induced membrane curvature in coarse-grained simulations. Biophys J 120, 3211–3221. 10.1016/j.bpj.2021.05.029

56. Gomez-Llobregat, J., Elias-Wolff, F., and Linden, M. (2016) Anisotropic Membrane Curvature Sensing by Amphipathic Peptides. Biophys J 110, 197–204. 10.1016/j.bpj.2015.11.3512

57. Pajtinka, P., and Vácha, R. (2024) Amphipathic Helices Can Sense Both Positive and Negative Curvatures of Lipid Membranes. The Journal of Physical Chemistry Letters 15, 175–179. 10.1021/acs.jpclett.3c02785

58. Paul, S., Audhya, A., and Cui, Q. (2024) Delineating the shape of COat Protein complex-II coated membrane bud. Pnas Nexus. ARTN pgae305 10.1093/pnasnexus/pgae305

59. Lincoff, J., Helsell, C. V. M., Marcoline, F. V., Natale, A. M., and Grabe, M. (2024) Membrane curvature sensing and symmetry breaking of the M2 proton channel from Influenza A. Elife. 10.7554/eLife.81571

60. Boesze-Battaglia, K., and Albert, A. D. (1992) Phospholipid distribution among bovine rod outer segment plasma membrane and disk membranes. Exp. Eye Res. 54, 821–823

61. Sander, C. L., Sears, A. E., Pinto, A. F. M., Choi, E. H., Kahremany, S., Gao, F., Salom, D., Jin, H., Pardon, E., Suh, S., Dong, Z., Steyaert, J., Saghatelian, A., Skowronska-Krawczyk, D., Kiser, P. D., and Palczewski, K. (2021) Nano-scale resolution of native retinal rod disk membranes reveals differences in lipid composition. J Cell Biol 220. 10.1083/jcb.202101063

62. Marsh, D. (2006) Elastic curvature constants of lipid monolayers and bilayers. Chem. Phys. Lipids 144, 146–159. 10.1016/j.chemphyslip.2006.08.004

63. Heath Jeffery, R. C., Thompson, J. A., Lo, J., Chelva, E. S., Armstrong, S., Pulido, J. S., Procopio, R., Vincent, A. L., Bianco, L., Battaglia Parodi, M., Ziccardi, L., Antonelli, G., Barbano, L., Marques, J. P., Geada, S., Carvalho, A. L., Tang, W. C., Chan, C. M., Boon, C. J. F., Hensman, J., Chen, T. C., Lin, C. Y., Chen, P. L., Vincent, A., Tumber, A., Heon, E., Grigg, J. R., Jamieson, R. V., Cornish, E. E., Nash, B. M., Borooah, S., Ayton, L. N., Britten-Jones, A. C., Edwards, T. L., Ruddle, J. B., Sharma, A., Porter, R. G., Lamey, T. M., McLaren, T. L., McLenachan, S., Roshandel, D., and Chen, F. K. (2024) Retinal Dystrophies Associated With Peripherin-2: Genetic Spectrum and Novel Clinical Observations in 241 Patients. Invest Ophthalmol Vis Sci. 10.1167/iovs.65.5.22

64. Cavanaugh, B. L., Milstein, M. L., Boucher, R. C., Tan, S. X., Hanna, M. W., Seidel, A., Frederiksen, R., Saunders, T. L., Sampath, A. P., Mitton, K. P., Zhang, D. Q., and Goldberg, A. F. X. (2024) A new mouse model for pattern dystrophy exhibits functional compensation prior and subsequent to retinal degeneration. Human Molecular Genetics 33, 1916–1928. 10.1093/hmg/ddae128

65. Sharma, Y. V., Cojucaru, R. I., Ritter, L. M., Khattree, N., Brooks, M., Scott, A., Swaroop, A., and Goldberg, A. F. X. (2012) Protective Gene Expression Changes Elicited by an Inherited Defect in Photoreceptor Structure. PLoS One 7, e31371. 10.1371/journal.pone0031371

66. Goldberg, A. F., Ritter, L. M., Khattree, N., Peachey, N. S., Fariss, R. N., Dang, L., Yu, M., and Bottrell, A. R. (2007) An intramembrane glutamic acid governs peripherin/rds function for photoreceptor disk morphogenesis. Invest. Ophthalmol. Vis. Sci. 48, 2975–2986. 48/7/2975 [pii];10.1167/iovs.07-0049 [doi]

67. Schagger, H., and Vonjagow, G. (1991) Blue Native Electrophoresis for Isolation of Membrane-Protein Complexes in Enzymatically Active Form. Anal. Biochem. 199, 223–231. Doi 10.1016/0003-2697(91)90094-A

68. Wittig, I., Braun, H. P., and Schägger, H. (2006) Blue native PAGE. Nature Protocols 1, 418–428. 10.1038/nprot.2006.62

69. Qi, Y., Ingolfsson, H. I., Cheng, X., Lee, J., Marrink, S. J., and Im, W. (2015) CHARMM-GUI Martini Maker for Coarse-Grained Simulations with the Martini Force Field. J Chem Theory Comput 11, 4486–4494. 10.1021/acs.jctc.5b00513

70. de Jong, D. H., Singh, G., Bennett, W. F., Arnarez, C., Wassenaar, T. A., Schafer, L. V., Periole, X., Tieleman, D. P., and Marrink, S. J. (2013) Improved Parameters for the Martini Coarse-Grained Protein Force Field. J Chem Theory Comput 9, 687–697. 10.1021/ct300646g

71. Abraham, M. J., Murtola, T., Schulz, R., Páll, S., Smith, J. C., Hess, B., and Lindahl, E. (2015) GROMACS: High performance molecular simulations through multi-level parallelism from laptops to supercomputers. SoftwareX 1**-****2**, 19–25. 10.1016/j.softx.2015.06.001

72. Hossein, A., and Sodt, A. J. (2023) MembraneAnalysis.jl: A Julia package for analyzing molecular dynamics simulations of lipid membranes. The Journal of Open Source Software. 10.21105/joss.05380

